# Ift43 Controls the Ciliary Levels of Gli2 and Gli3

**DOI:** 10.64898/2026.01.14.699321

**Authors:** Michael W. Stuck, Mohona Gupta, Luke N. Knutson, Paurav B. Desai, Karyn L. Robert, Jeffrey J. Anuszczyk, Abigail O Smith, Delayna Paulson, Sumeda Nandadasa, William Devine, Cecilia W Lo, Timothy C. Cox, Darci M. Fink, Gregory J. Pazour

## Abstract

Intraflagellar transport (IFT) drives the bidirectional movement of trains composed of IFT-A, IFT-B, and BBSome complexes that build and maintain cilia while supporting their signaling functions. Over evolution, IFT became integral to Hedgehog signaling by directing the dynamic movements of receptors and Gli transcription factors that fine-tune pathway output. The IFT-A complex contains six subunits, but the smallest, Ift43, remains poorly characterized and is absent from many ciliated species, suggesting specialized roles in signaling rather than core ciliogenesis. Here we show that loss of *Ift43* in mice causes mid-gestation lethality with severe craniofacial defects, exencephaly, abdominal wall defects with exposed viscera, edema, and limb patterning defects. At the cellular level, Ift43 deficiency reduces both the number and length of cilia and blocks induction of Gli1 following pathway activation by the agonist SAG. Although Smoothened relocalizes to cilia normally, *Ift43* mutants abnormally accumulate Gli2 and Gli3 at ciliary tips before stimulation and continue to generate repressor forms after activation. Conversely, Ift43 overexpression increases basal Gli2 cleavage, revealing an unanticipated role for Ift43 in regulating Gli processing. Together, these findings identify Ift43 as a key IFT-A component that links ciliary assembly to Hedgehog signal transduction and helps set the balance between Gli activator and repressor forms.

## Introduction

In eukaryotes, cilia and flagella play crucial roles in development and maintenance of health by serving as sensory organelles and by producing force. The sensory functions underlie our senses of sight and smell and are critical during development and in tissue homeostasis. Ciliary motility moves fluid over epithelia in the respiratory and reproductive tracts and in the ventricles of the brain, as well as propel sperm for fertilization. In development, cilia play critical roles in breaking left-right symmetry, which gives rise to the asymmetric placement of our thoracic and abdominal organs and is critical to development of a heart that can support systemic and pulmonary blood flow. In addition, cilia are integral to Hedgehog signaling as the pathway is organized in the cilium. Activation of Hedgehog signaling initiates with the binding of a Hedgehog ligand to patched1 (Ptch1) located in the ciliary membrane. This binding causes Ptch1 to exit cilia (Rohatgi et al., 2007) and relieves its inhibition of the pathway, allowing the seven transmembrane receptor smoothened (Smo) to accumulate in cilia and become activated (Corbit et al., 2005). The activation involves the accumulation of the Gli transcription factors at the ciliary tip where they are converted to transcriptional activators before moving to the nucleus to activate transcription (Haycraft et al., 2005). Intraflagellar transport (IFT) is tightly connected to Hedgehog signaling, with IFT defects causing attenuation of signaling (Huangfu et al., 2003) and disruption of the dynamic redistribution of components that normally occurs during signaling (Keady et al., 2012).

During IFT, kinesin-2 (Cole et al., 1998) and dynein-2 (Pazour et al., 1998) motors transport large multimeric IFT trains from the cell body into and along the ciliary microtubules in the anterograde and retrograde directions. In addition to roles in Hedgehog signaling, this process drives ciliary assembly in most eukaryotes (Rosenbaum and Witman, 2002). The IFT trains are composed of three conserved subcomplexes, IFT-A, IFT-B, and the BBSome. In this work, we focus on Ift43, the smallest subunit of IFT-A (Cole, 2003). In cryo IFT-A structures (Hesketh et al., 2022; Jiang et al., 2023; Ma et al., 2023; Meleppattu et al., 2022), Ift43 amino acids 128-184 of this 206 residue protein (numbered from mouse sequence) are embedded into the IFT-A particle and make extensive contacts with Ift121 and Ift139. In the vertebrate structure, the N-terminal 127 residues and the C-terminal 22 residues are not observed and are suggested to be unstructured by AlphaFold. Four human patient variants have been identified. Two variants disrupt the start codon: c.A1G and c.T2A, which remove the first 22 residues if translation initiates at the next start codon. Patients homozygous for these variants develop skeletal dysplasias (Arts et al., 2011; Duran et al., 2017). Another family with a skeletal dysplasia carried a W179R variant (Duran et al., 2017). W179 (W172 in mouse) is located in the interface with Ift121 and appears to destabilize the IFT-A complex (Jiang et al., 2023). A fourth family carrying E34K mutations developed isolated retinal degeneration (Biswas et al., 2017). E34 (also E34 in mouse) is within the disordered region at the N-terminus.

In this work, we show that Ift43 is critical to mouse development, with null embryos showing severe craniofacial abnormalities along with significant defects in the heart, skeleton, lymphatic system, and other organs. *Ift43* null fibroblasts show a reduced number of shorter cilia that are not capable of Hedgehog signaling. The mutants inappropriately accumulate Gli2 and Gli3 at the ciliary tip under basal conditions. At basal conditions, Gli3, and Gli2 to a lesser extent, are normally processed into cleaved repressor forms. Ift43 mutants continue to process Gli3 and Gli2 into the repressor forms after pathway activation and overexpression of Ift43 promotes the production of cleaved Gli2 beyond what is normally seen. These findings suggest that Ift43 is crucial for the transport and processing of Gli2 and Gli3, a role not been previously appreciated.

## Results

### Conservation of Ift43 across eukaryotes: Ift43 is similar to Ift27

In 2013, van Dam and colleagues examined the distribution of IFT proteins across a wide spectrum of ciliated and non-ciliated eukaryotes. As expected, IFT proteins were highly conserved in ciliated organisms, but outliers were observed (van Dam et al., 2013). In IFT-A, Ift43 was the least conserved subunit and was missing from numerous organisms that retained the rest of IFT-A. Curiously, Ift43 was often missing from organisms that lacked Ift25 or Ift27. This distribution is intriguing as mouse Ift25 and Ift27 are required for Hedgehog signaling but not ciliary assembly (Eguether et al., 2014; Keady et al., 2012), suggesting that Ift43 also could have specialized functions. Genome assemblies have matured since 2013, and so we re-examined the genomes using reiterative BLAST searches followed by phylogenetic analysis to identify Ift43 along with Ift25, Ift27, and other IFT components (Fig 1). While numerous differences were observed in our data set as compared to van Dam, the overall trends remained. Ift25 and Ift27 are typically lost together, but several organisms (*Ciona*, *Monosiga*, *Emiliania*, *Paramecium*, *Tetrahymena*) lost Ift25 while retaining Ift27. There is a strong, but not perfect, correlation between the presence of Ift27 and Ift43, with most organisms either retaining both proteins or losing both proteins. Exceptions include *Monosiga*, *Emiliana*, and *Trichomonas* that have Ift27 but lack Ift43. Conversely, Ift43 is found in *Caenorhabditis* and *Drosophila* where Ift25 and Ift27 are missing. Curiously, gene loss in *Anopheles* mosquitos varies with Ift43 being found in some species but not others.

**Figure 1.**
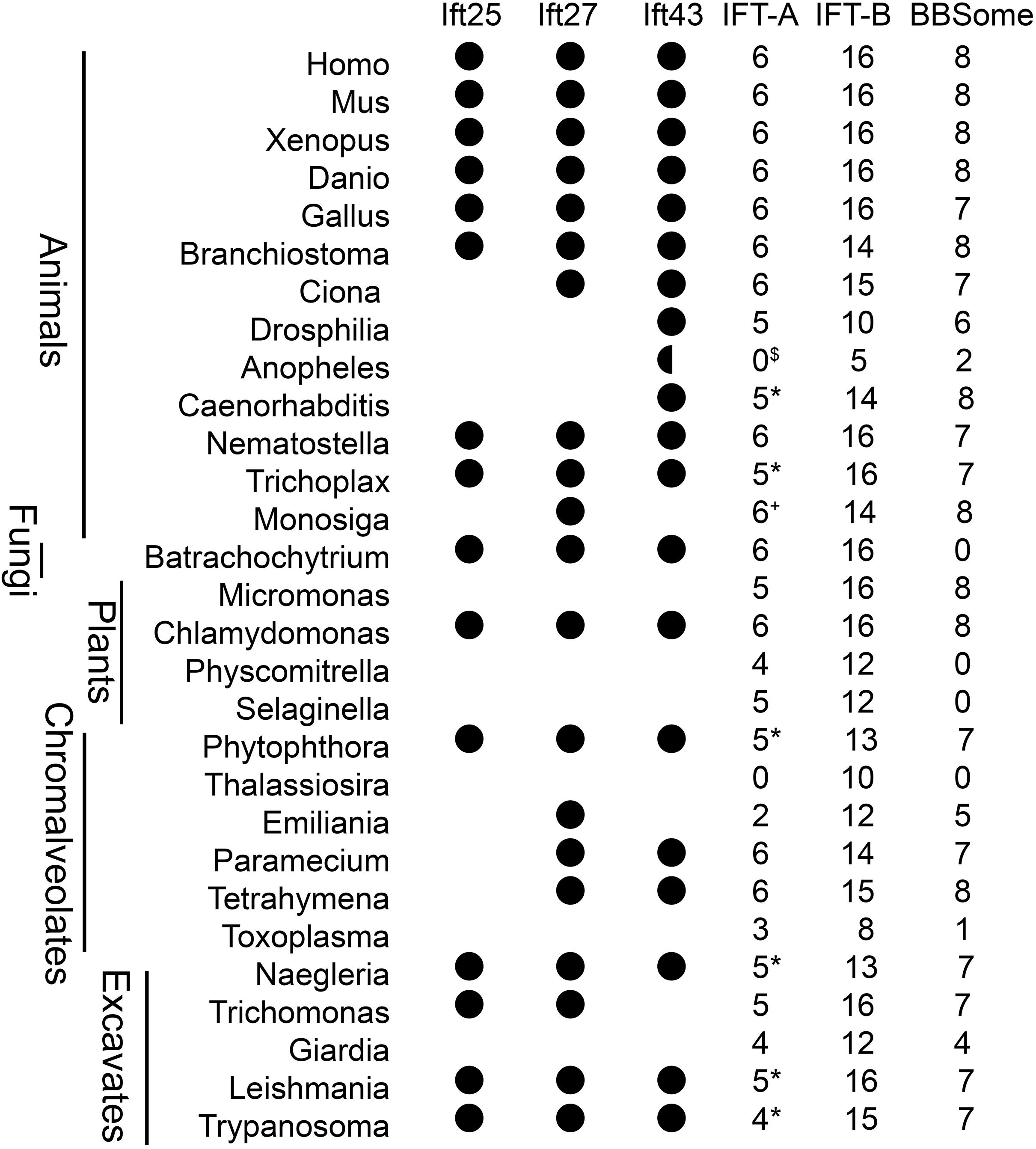
Conservation of Ift43. The presence of *Ift25*, *Ift27* and *Ift43* in an organism is denoted by a black circle. A half circle in *Anopheles* indicates that it is present in some mosquito species but not in all. Numbers under IFT-A, IFT-B, and BBSome indicate how many of the 6, 16, and 8 core proteins were identified in the species in the work of (Dobbelaere et al., 2023). *Ift43 was not found in this large screen but found in our analysis. ^+^Ift43 was found in the large screen but not found in our analysis. ^$^data is for *Anopheles gambiae*; retention of IFT genes varied by *Anopheles* species. Data for humans, chickens (*Gallus*) and mosquitoes (*Anopheles*) were obtained from NCBI Gene. See S1 Data for data to support this figure.

Overall, the level of sequence identity in Ift43 is low across eukaryotes (S1 Data), which makes the identification of Ift43 challenging. Thus, it is possible that some Ift43-like sequences represent other proteins. Within the animals, *Caenorhabditis* Ift43 has low sequence identity to other members of the family. However, *Caenorhabditis* Ift43 is likely a true IFT-A subunit as a tagged version localized to amphid and phasmid cilia in transgenic worms and immunoprecipitation of the IFT-A subunit Ift140 co-precipitated this protein (Yi et al., 2017). Human (Mukhopadhyay et al., 2010) and *Chlamydomonas* Ift43 (Behal et al., 2012) purified with IFT-A. *Leishmania* (Meleppattu et al., 2022), human (Hesketh et al., 2022; Jiang et al., 2023) and *Tetrahymena* (Ma et al., 2023) Ift43 were recently shown to be part of the IFT-A structure, but the evidence that the other proteins are true IFT-A subunits remains to be determined.

### Mouse Ift43 is required for embryonic development

To understand the function of Ift43 in mouse development, we obtained the *Ift43^tm1a^* targeted allele (Skarnes et al., 2011). The *Ift43^tm1a^* mouse carries a promotor-less beta-galactosidase gene trap cassette and a neomycin selectable marker in *Ift43* intron two and has exon three flanked by Flox sites (S2 Data) The gene trap insertion is expected to prevent production of the normal mRNA, thus creating a null or strong hypomorphic allele. *Ift43^tm1a^* heterozygotes have no obvious phenotypes and breed normally. However, no homozygous offspring were obtained from intercrosses of heterozygotes (27 *Ift43^+/+^*, 56 *Ift43^tm1a/+^* offspring from 16 litters, Chi-square p<0.0001 with an expected distribution of 1:2:1). To determine the time of embryo death, pregnant females from timed matings were dissected at embryonic (E) days 11.5, 12.5, 13.5 and 14.5. Embryos at all time points showed strong craniofacial abnormalities and malformed hearts. At E14.5, several of the affected were dead, and the living ones had only sporadic heartbeats, suggesting they were close to demise (S1 Table).

The *Ift43^tm1a^* allele was converted to *Ift43^tm1b^*and *Ift43^tm1e^* (flox) alleles by crossing to germline Zp3-Cre and FlpO, respectively (S2 Data). *Ift43^tm1b^* retains the promotor-less beta-galactosidase gene trap cassette in exon 2 but is missing the neomycin selection marker and exon 3. It is expected that this arrangement will produce a null allele, as the genetrap should capture any transcription that starts from the promoter. If any transcripts are made and processed, exon 4 is expected to be spliced onto exon 2, resulting in a frameshift and termination after the initial 44 codons (S2 Data). *Ift43^tm1b^* heterozygotes bred normally but no homozygous offspring were obtained at weaning (40 *Ift43^+/+^*, 52 *Ift43^tm1b/+^* offspring from 19 litters, Chi-square p<0.0001 with an expected distribution of 1:2:1). The number of heterozygotes was slightly less than expected (Chi-square p=0.047 if the expected distribution is 1:2:0). The recovered heterozygotes obtained appeared largely normal, although a few minor phenotypes were observed (S4 Data). Dissection of pregnant females obtained from timed mating of heterozygotes showed phenotypes very similar to the *Ift43^tm1a^*homozygotes, with most E14.5 mutant embryos alive but with weak sporadic heartbeats. At E15.5, about ½ of mutant embryos were dead, but embryos with detectable heartbeats were able to be collected for phenotyping (S1 Table). Homozygous *Ift43^flox/flox^* mice showed no obvious phenotypes and bred normally.

### Ift43 loss causes extensive structural birth defects

All *Ift43^tm1b/tm1b^* mutants showed extensive structural birth defects (Fig 2) including severe craniofacial abnormalities with a gaping mouth and marked exencephaly. In addition, all mutants had extensive fluid accumulation under the skin (hydrops) of the body, and the heart and liver were exposed due to an abdominal wall closure defect. A variety of digit defects, including oligodactyly and polydactyly were observed. Intriguingly, a subset of mutants showed a curled tail, a phenotype commonly associated with defective convergent extension caused by the disruption of the planar cell polarity pathway (Ybot-Gonzalez et al., 2007). The heterozygotes visually appeared normal.

**Figure 2.**
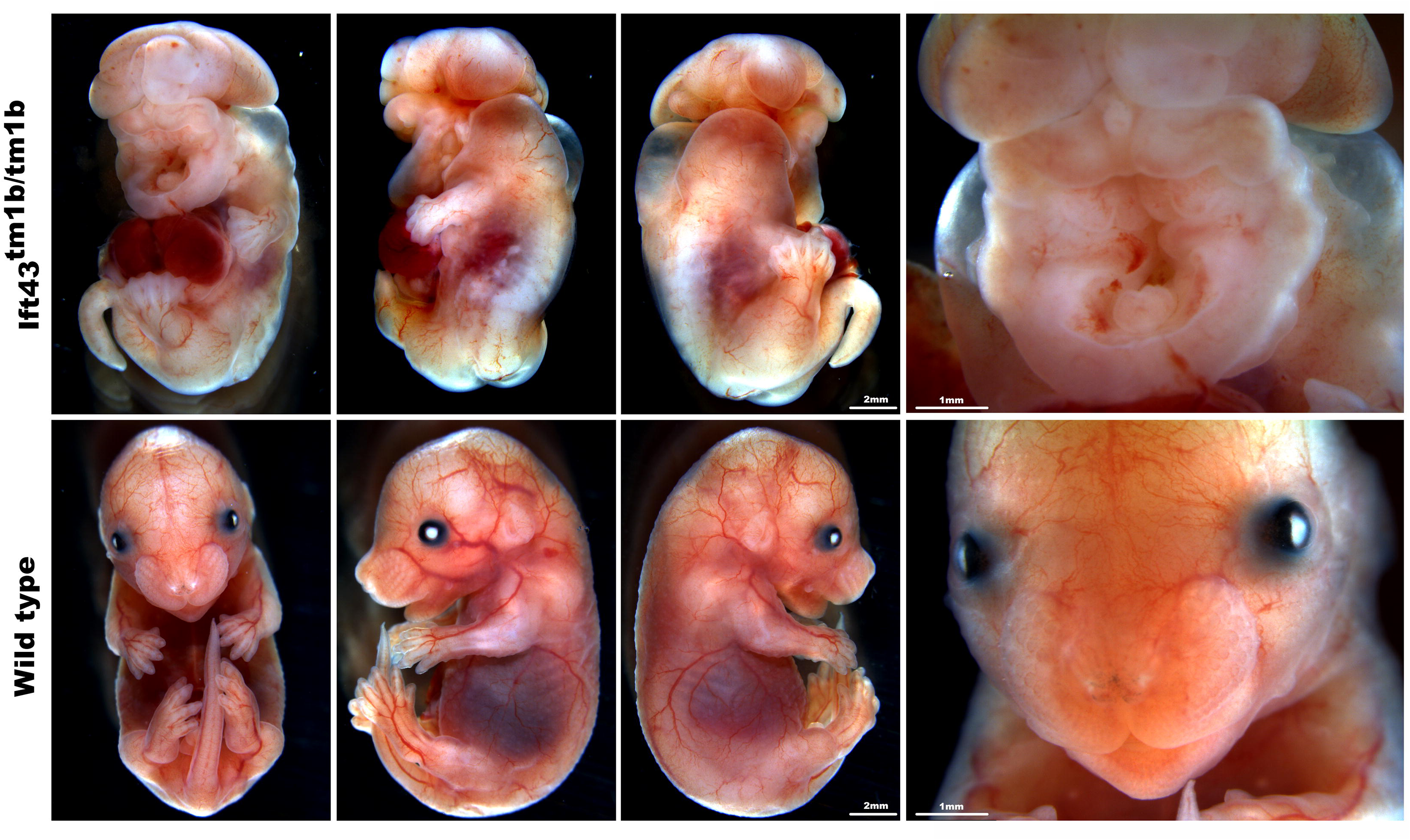
Loss of Ift43 causes structural birth defects. Light microscopy images of E15.5 embryos. Scale bar = 2 mm for whole body images and 1 mm for facial images.

Mutant and control embryos were subjected to necropsy, computed tomography (CT) scans, episcopic fluorescence image capture (EFIC), and optical projection tomography (OPT) to characterize the structural birth defects (Figs 3-5, S1-S2 Figs, S4 Data). Necropsy documented the complete penetrance of excencephaly and a variety of digit defects including oligodactyly due to the loss of digit five and polydactyly due to duplications of digits one, three, or five (S4 Data). CT, EFIC, and OPT imaging showed that the exencephaly was accompanied by extensive brain malformations. In some embryos, the right and left lobes were fused (holoprosencephaly), while in others there was an extra medial telencephalic lobe (Fig 3A-3D, 3N-3O). EFIC showed extensive organ defects including cardiac abnormalities that will be discussed subsequently. The mutants typically had a diaphragmatic hernia, so the abdominal and thoracic organs were intermingled with the liver projecting up and around the heart and lungs. The pulmonary organ appeared to consist of central hypoplastic single lung or two fused hypoplastic lungs that extended into the center of the liver (Fig 3E-3H). The spinal column was malformed and wavy (Fig 3L-3M) with abnormal vertebrate fusion (spinal dysraphism) (Fig 3J-3K). This was confirmed in the OPT imaging, which also revealed significant morphological anomalies of the somites, particularly in the tail (S2 Fig). Stomach position was variable with examples of left-, center-, and right-sided placement observed in the mutants (Fig 3I).

**Figure 3.**
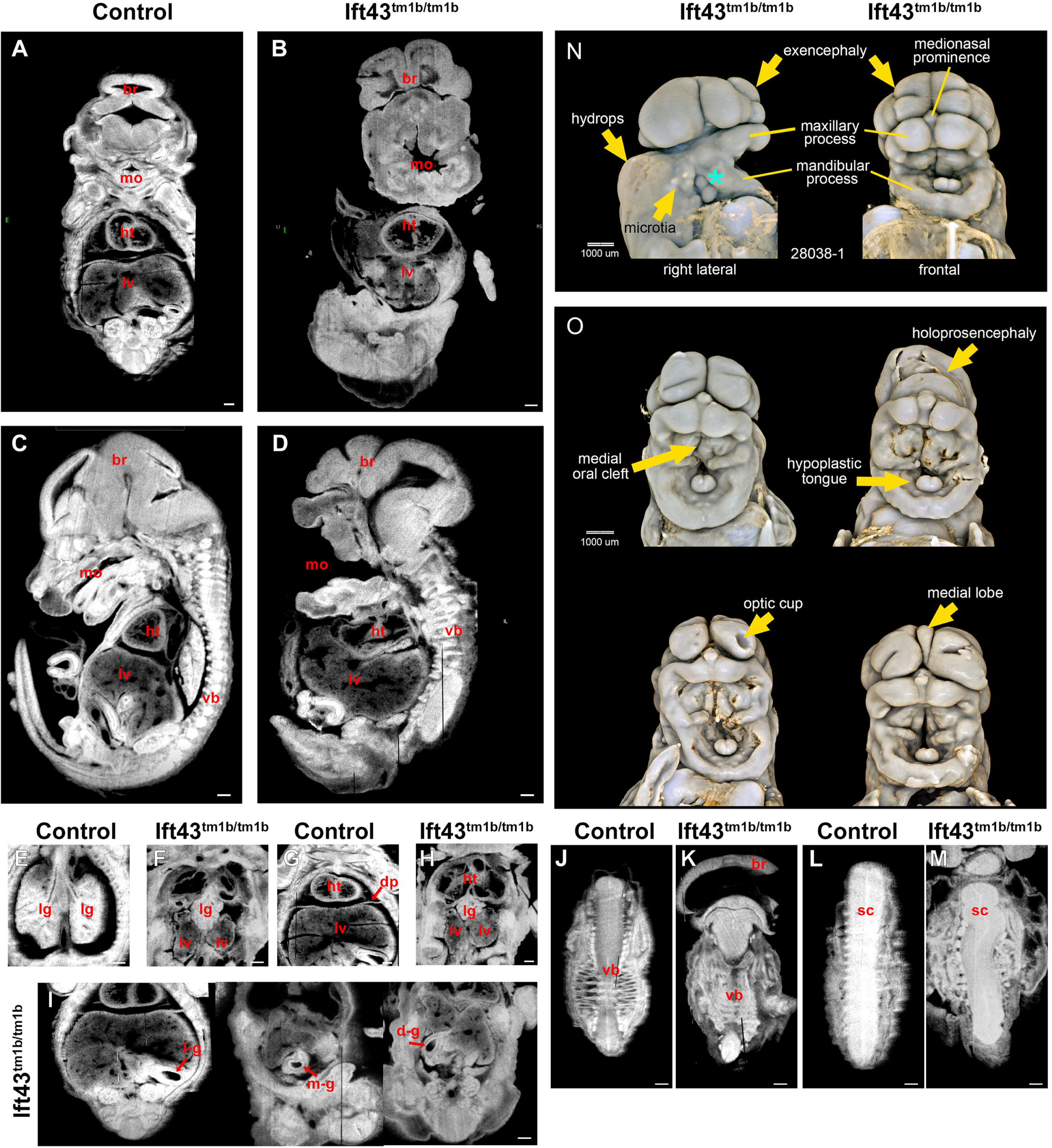
EFIC and OPT imaging reveals extensive birth defects at E15.5. EFIC (**A-M**) and OPT (**N-O**) images of E15.5 embryos. Controls are *Ift43^+/+^* or *Ift43^tm1b/+^* and mutants are *Ift43^tm1b/tm1b^*. **(A-D**) *Ift43* mutant embryos have an abdominal-chest wall defect with protruding bowel, liver (lv) and heart (ht), along with vertebral body (vb) abnormalities, brain (br) malformations, severe craniofacial defects including a gaping mouth (mo). Scale bars = 1mm. **(E-H**) *Ift43* mutant embryos have diaphragmatic hernia (dp), allowing the liver (lv) to extend into the thoracic cavity where it surrounds a single lobed abnormal lung (lg) below the heart (ht). Scale bars = 1 mm. **(I**) Situs anomalies involving the stomach were observed, with one mutant exhibiting midline stomach or mesogastria (m-g), another showing right-sided stomach (dextrogastria, d-g), and one showing normal left-sided stomach (levogastria, l-g). Scale bars = 1 mm. **(J-M**) Images showing defective vertebrate (vb) and spinal cord (sc) in the mutant animals. Brain (br) in (**K)** shows evidence of holoprosencephaly. Scale bars = 1 mm. **(M-N**) OPT images of the heads of experimental E15.5 embryos (*Ift43^tm1b/tm1b^*). Scale bar = 1 mm. See S4 Data for necropsy, CT, and EFIC phenotyping.

### Ift43 mutants have severe cranial facial abnormalities

To more carefully characterize the craniofacial abnormalities, six E15.5 *Ift43^tm1b/tm1b^* homozygous mutant embryos along with controls (*Ift43^+/+^* or *Ift43^tm1b/+^*) were examined by high-resolution three-dimensional OPT imaging (Fig 3N-3O, S1 and S2 Figs). This assessment revealed that, while the maxillary and mandibular processes were present and particularly prominent for the latter, the frontonasal prominence-derived tissue was severely deficient. There was minimal cranial tissue supporting the brain. For most mutants, there was no evidence of any definitive ocular tissue. However, one mutant exhibited a round depression on the left telencephalon, suggesting initiation of optic cup invagination. All other mutants exhibited a single fold on each side of the telencephalon, which may represent failed optic cup invagination (Fig 3O). Despite the comparatively large mandibular tissue, the tongue was severely hypoplastic in all homozygotes, yet papillae were evident, suggesting normal differentiation of remaining tissue. Assessment of the reconstructed serial sections from OPT imaging indicates that the mandibular processes may not have merged at the midline (S2 i’ Fig). All homozygotes showed a medial intraoral cleft, which may reflect a separation of midline cranial tissues rather than true cleft palate, since the clefts extended almost to the base of the midbrain (S2 iii’ Fig). An overt medial cleft lip was also observed in two embryos, one of which had holoprosencephaly. There was asymmetry in the appearance of the dorsal intraoral tissue in 6 of the 7 homozygotes imaged, with 5 showing smaller right-sided tissue (Fig 3N-3O). External ear (pinna) development was very abnormal, with all mutant auricular tissue resembling human grade III microtia (Fig 3N). Two large tissue masses were also evident at the proximal (ramus) end of the mandible, between the mandible proper and the remnant auricular tissue (cyan asterisk in Fig 3N). The origin of these masses is unknown, but they may be derivatives of the Hillocks of His. On one or both sides of most homozygotes, there was a likely fluid-filled ‘blister’ around the “hinge” region between the mandible and maxilla (S2 ii’ Fig).

### Ift43 mutants have extensive cardiac malformations

Collection of embryos from timed matings showed that at E15.5 about half of the mutant embryos were dead and the remaining living animals showed only sporadic heart beats indicating that they were close to death (S1 Table). At this age, heart development is largely complete with a functional four chambered heart. To understand the importance of Ift43 in heart development, embryos at E15.5 were examined by EFIC imaging. In contrast to the controls with septa separating the left and right atria (Fig 4A-4B), mutants showed a common atrium with a common atrioventricular valve (Fig4C). Mutants showed evidence that the truncus arteriosus failed to separate into the pulmonary artery and aorta causing persistent truncus arteriosus (Fig 4B, 4D). In addition, mutants were observed to have dextroversion or rightward rotation of the heart (Fig 4C, 4F) and superior-inferior ventricles where the ventricles are positioned one above the other, rather than side-by-side (Fig 4C, 4F). In one case, this was accompanied by a muscular ventricular septal defect (Fig 4F).

**Figure 4.**
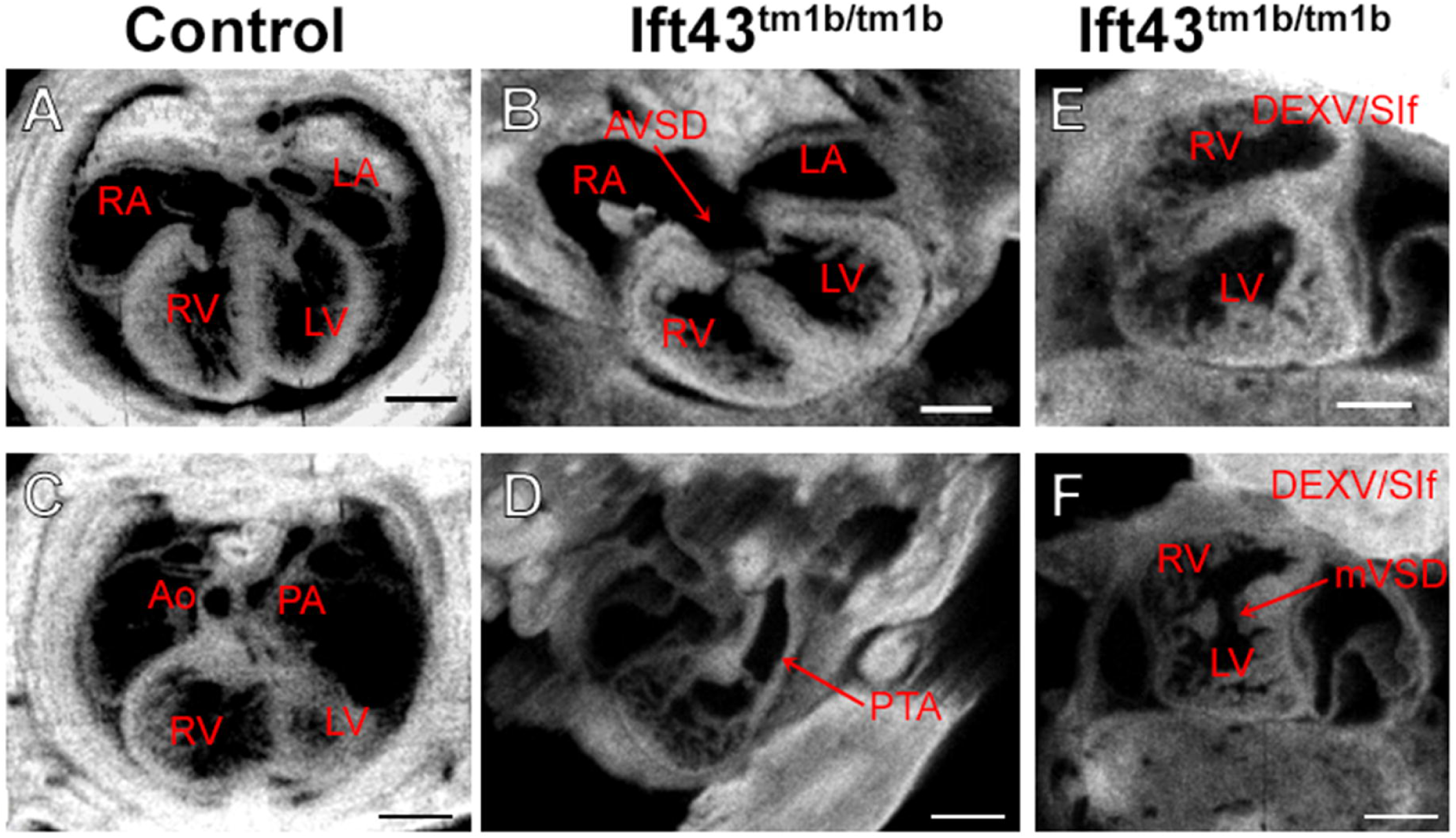
Ift43 KO mouse embryos show disturbances of laterality and other structural birth defects. **(A**) Wild-type heart showing normal four chamber heart comprising right (RA) and left atria (LA) and right (RV) and left ventricles (LV) with normal intact ventricular septum. Scale bar = 1.5 mm. **(B**) Four chamber view of a mutant embryo showing an atrioventricular septal defect (AVSD) and its common atrioventricular valve. Scale bar = 1.5 mm. **(C-D**) Great arteries in a normal mouse embryo (**C**), while in a mutant (**D**), a solitary arterial trunk (arrow) indicates possible persistent truncus arteriosus (PTA). Scale bar = 1.5 mm. **(E)** Mutant showing right-sided heart with dextroversion (DEXV) and superior-inferior (human orientation) (SIf) ventricles. Scale bar = 1.5 mm. **(F)** A right-sided heart with dextroversion, superior-inferior ventricles, and a muscular ventricular septal defect (mVSD, arrow). Scale bar = 1.5 mm. See S4 Data for necropsy, CT, and EFIC phenotyping.

### Ift43 knockout impairs skin lymphatic vessel patterning during embryonic development

Hydrops, the accumulation of fluid under the skin, suggests lymphatic defects. Lymphatic endothelial cells are ciliated and disrupting these cilia via an *Ift20* knockout caused edema that was associated with increased and more variable lymphatic vessel caliber and branching, as well as aberrant red blood cell infiltration. (Paulson et al., 2021). To determine if the loss of *Ift43* would dysregulate lymphatic plexus formation, we examined whole mount E15.5 dorsal skin from *Ift43^tm1b/tm1b^* knockouts and littermate controls. Staining skin with antibodies against acetylated α-tubulin and Arl13b detected abundant cilia in controls. In contrast, no cilia-like structures were stained in knockouts with Arl13b staining (S3 Fig).

However, approximately normal numbers of short cilia were observed when tissue was stained with acetylated α-tubulin antibodies (Fig 5G-5J), suggesting that Ift43 is required for delivery of Arl13b to cilia. Similar to what we observed in *Ift20* knockouts, lymphatic vessel density was higher in *Ift43* knockouts (Fig 5A, 5D) and vessel diameter was increased and more variable in *Ift43* knockouts (Fig5A-C), with some diameters measuring nearly 400 μm (Fig 5B). No significant difference in proliferation was detected by phospho-histone H3 staining of Prox1-positive lymphatic endothelial cells at this point in development (Fig 5E, 5F and S4 Fig). These results indicate that primary cilia on lymphatic endothelial cells restrict excessive lymphangiogenesis and promote appropriate lymphatic vessel network formation.

**Figure 5.**
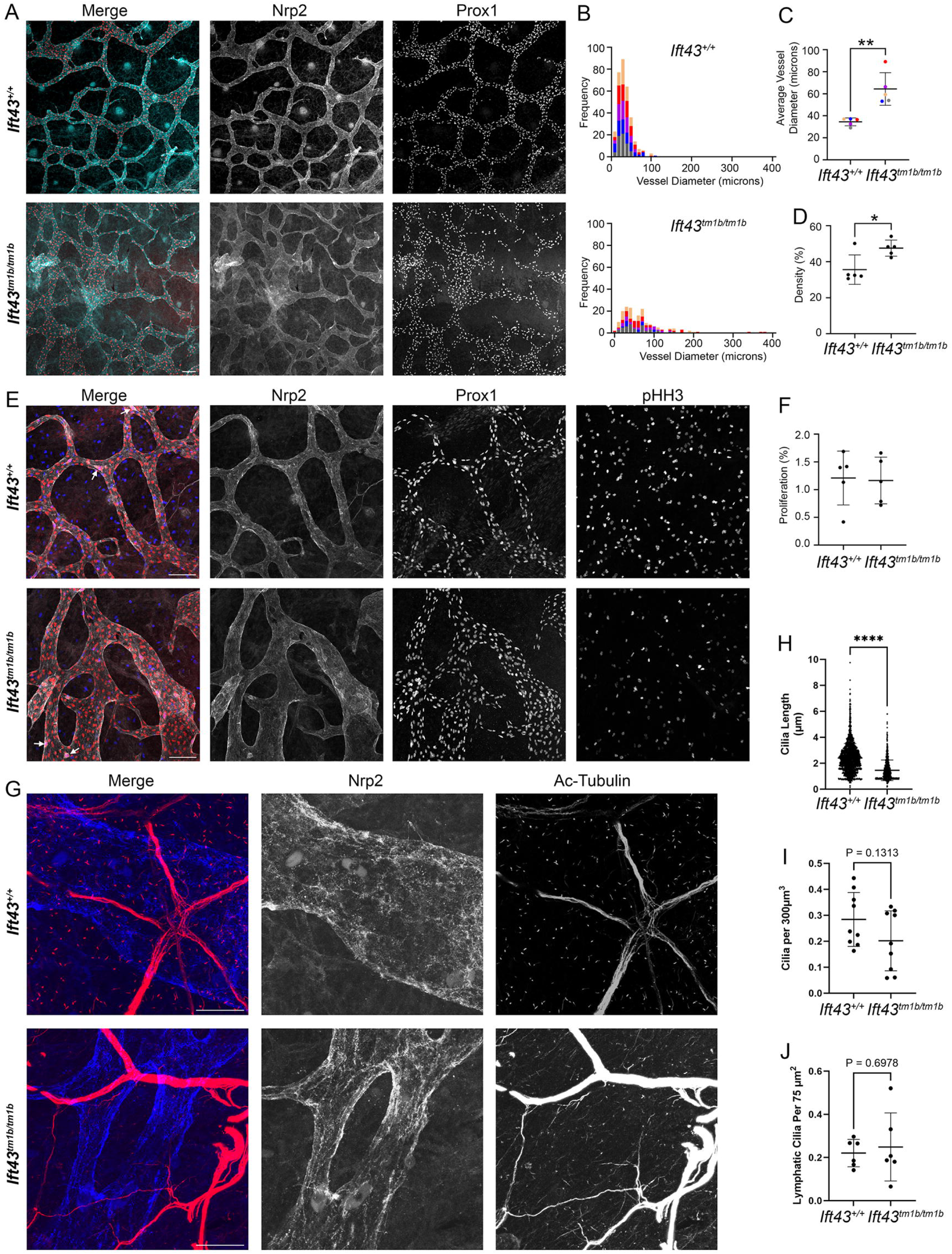
Ift43 loss affects embryonic lymphatic vessel development. **(A)** Immunofluorescence staining for lymphatic vessel networks (Nrp2, green; Prox1, red) in whole mount E15.5 dorsal skin. Images are maximum intensity z-projections of images taken at 1.0 µm intervals. Scale bar = 100 µm. **(B)** Frequency distributions of vessel diameter. **(C)** Quantification of average lymphatic vessel diameter. Data represents 18 fields of view across 5 control and 5 experimental embryos across 5 litters. Each dot is the average from one embryo. **p = 0.0092 by unpaired t-tests with Welch’s correction. **(D)** Quantification of percent area filled with lymphatic vessel signal. Each dot is the average from one embryo. *p = 0.0275 by unpaired t-tests with Welch’s correction. **(E)** Immunofluorescence staining of whole mount E15.5 dorsal skin to measure proliferation. Lymphatic vessels labeled via Nrp2 and Prox1. Proliferating cells labeled via phospho-histone H3 (pHH3). White arrows highlight colocalization of Prox1 and pHH3, indicating proliferating lymphatic endothelial cells. Images are maximum projections of stacks taken at 1.0 µm intervals. Scale bar = 100 µm. **(F)** Quantification of percent lymphatic endothelial cell nuclei undergoing proliferation. Data represents 19 fields of view across 5 control and 5 experimental embryos across 5 litters. Each dot is the average from one embryo. See Fig S3 for details on quantification. **(G)** Immunofluorescence staining for primary cilia and lymphatic vessels in whole mount E15.5 dorsal skin. Lymphatic vessel surface labeled via neuropilin-2 (Nrp2), lymphatic endothelial cell nuclei labeled via transcription factor Prox1, and primary cilia labeled via Arl13b. Images are maximum projections of stacks taken at 0.5 µm intervals. Scale bar = 20 µm. See Fig S2 for staining with Arl13b. **(H)** Quantification of cilia length of total cilia. Data represents 18 fields of view across 3 control and 3 mutant embryos across 3 litters. Each dot represents the length of one cilium. n = 2053 (control) and 956 (Ift43 mutant), ****p = <0.0001 by unpaired t-tests with Welch’s correction. **(I)** Quantification of total cilia per 300 µm^2^. Data represents 18 fields of view across 3 control and 3 mutant embryos across 3 litters. Each dot represents a field of view. p = 0.1 by unpaired t-tests with Welch’s correction. **(J)** Quantification of cilia on lymphatic endothelial cells per 75 µm^2^. Data represents 12 fields of view across 2 control and 2 KO embryos across 2 litters. Each dot represents one field of view. p = 0.7 by unpaired t-tests with Welch’s correction. Raw data underling the graphs are available in S5 Data.

### Ift43 is required for photoreceptor maintenance

To explore the role of Ift43 in photoreceptors, we deleted the floxed allele of *Ift43* using *iCre75*. This driver uses the rod opsin promoter to selectively express Cre recombinase in rod photoreceptor cells starting at about postnatal day 7 (P7) (Li et al., 2005). The *Ift43* floxed allele contains loxP sites flanking exon 3. Deletion of this exon is likely to produce a null allele due to the out-of-frame fusion of exons 2 and 4, which would allow for translation of only the first 44 residues before going out of frame. Outer segment elongation starts at about P9/P10 (LaVail, 1973; Obata and Usukura, 1992; Pearring et al., 2013). Due to the time required for the gene to be deleted and gene product decay, we would expect pathology to start during the elongation phase of outer segment development.

At P30, a point at which outer segments should be fully elongated, electroretinography (ERG) showed dysfunction in both the initial rod photoreceptor response to light (scotopic a wave) and the downstream responses (scotopic b wave) in experimental animals compared to controls. Cone-driven responses (photopic b wave) were not significantly affected at these points (Fig 6A). This is expected as we deleted the *Ift43* gene in rod photoreceptors, and the time points examined are too early for secondary cone loss. Electron microscopy of experimental P21 and P30 retinas showed significant disruption of the outer segments, with the outer segments irregularly shaped and shortened and the remaining discs being abnormal in size and orientation, with the phenotype more extensive at P30 (Fig 6Ba-f). Tannic acid and uranyl acetate label the membranes of newly forming discs exposed to the extracellular environment more darkly than the membranes of fully formed discs enclosed inside the outer segment. As expected, this method darkened the base of the photoreceptor outer segments in the control retinas. In the experimental animals, at P21, there was mixture of darkened and undarkened discs at the base of the outer segments. By P30, few outer segment bases were darkened in the experimental retinas, indicating that new discs were not being formed in the absence of Ift43 (Fig 6Bj-l). In addition, the inner segments of the experimental animals showed increased accumulation of extracellular vesicles as compared to controls (Fig 6Bg-i, 6C). The Golgi complex was mildly affected with occasional large vesicles observed within the structure (Fig 6C). At P25, significant rhodopsin was mislocalized in the inner segment and outer nuclear layer (Fig 6Da). Wheat germ agglutinin (WGA), which labels rhodopsin and other glycoproteins, also showed increased stain in the inner segment and outer nuclear layer (Fig 6Dc). As expected, cone opsins were not affected by the loss of Ift43 from rods (Fig 6Db). Although rhodopsin was mislocalized, Prom1 and Rom1 — outer segment-enriched proteins involved in disc morphogenesis and structural maintenance — and Stx3, an inner segment SNARE protein essential for vesicular trafficking to the outer segment, maintained normal distributions (S5 Fig). Notably, these proteins are often mislocalized in various forms of retinal degeneration, suggesting that the preservation of their localization in this model reflects a selective defect in rhodopsin trafficking rather than generalized disruption of outer or inner segment structure. These data show that Ift43 is important for the elongation and maintenance of the outer segment and are consistent with the finding that patients with *Ift43* variants develop retinal degeneration (Biswas et al., 2017).

**Figure 6:**
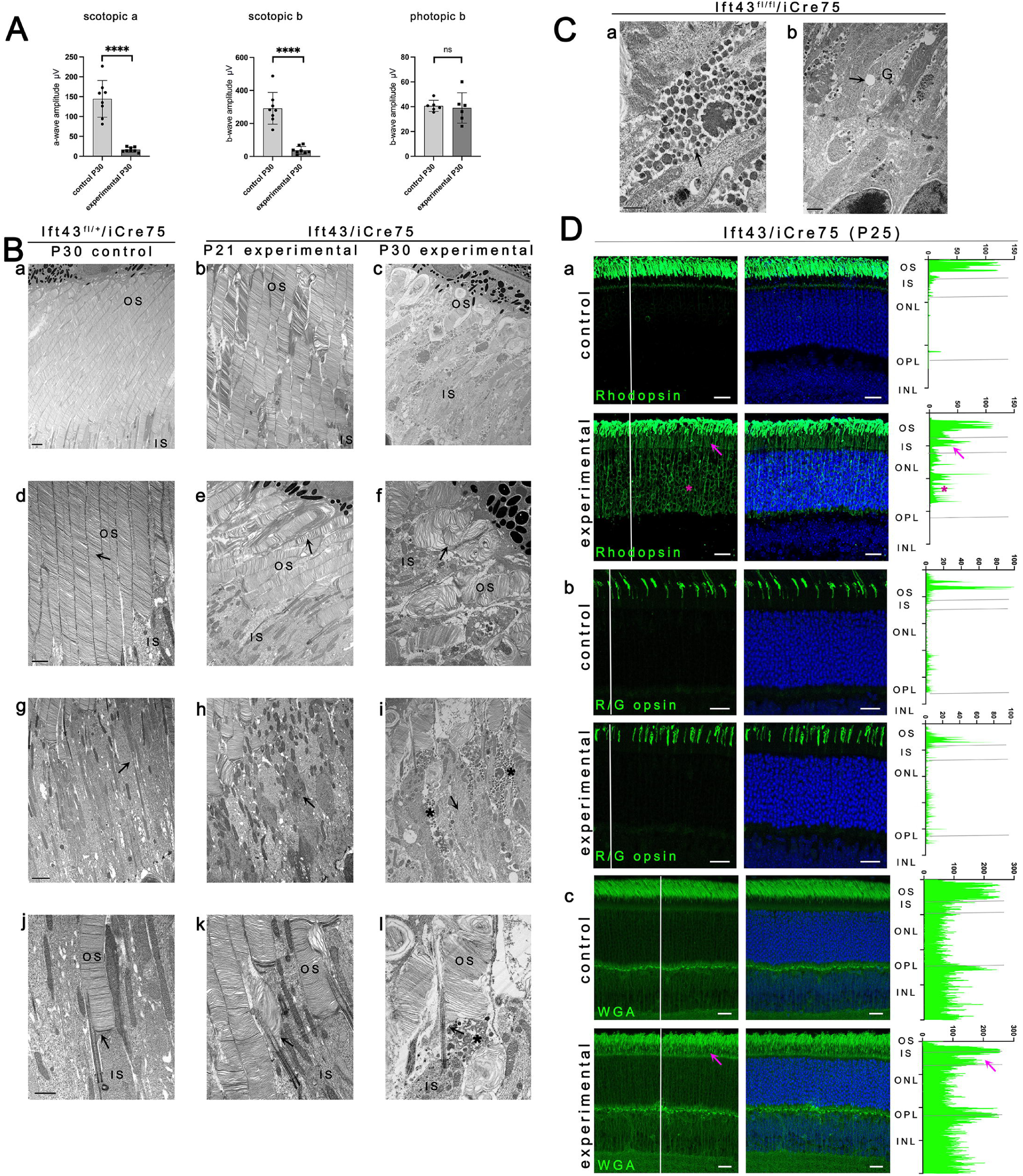
Ift43 loss induces photoreceptor degeneration and extracellular vesicle release from inner segment. **(A)** Electroretinograms (ERGs) of control (*Ift43^flox/+^/iCre75)* and experimental (*Ift43^flox/flox^/iCre75)* littermates at postnatal day 30 (P30). Each dot represents the ERG response from each eye of an animal. ****p<0.0001, ns not significant by unpaired t-tests. **(B)** Transmission electron microscopy images of retinal sections from control (*Ift43^flox/+^/iCre75)* and experimental (*Ift43^flox/flox^/ iCre75)* littermates at postnatal day 21 and 30 (P21 and P30). Scale bar = 2 µm (**Ba-Bc**), 1 µm (**Bd-Bf**), 2 µm (**Bg-Bi**), 1 µm (**Bj-Bl**). Arrows in **Bd-f** mark outer segments. Arrows in **Bg-i** mark inner segments and asterisks in **Bi** mark extracellular vesicles. Arrows in **Bj-k** mark examples of outer segment bases. Asterisk in **Bl** marks extracellular vesicles. OS: outer segment; IS: inner segment. n=3 animals examined by TEM. **(C)** High magnification TEM images of extracellular vesicles in the inner segment (arrow, **Ca**) and vesicle (arrow, **Cb**) in the Golgi complex (G) of experimental (*Ift43^flox/flox^/iCre75)* littermates at postnatal day 30. Scale bar = 0.5 μm (**Ca**), 1 μm (**Cb**). n=3 animals examined by TEM. **(D)** Confocal images of retinal cross sections (agarose embedded) at P25. Control (*Ift43^flox/+^/iCre75)* and experimental (*Ift43^flox/flox^/iCre75)* littermate mice stained with antibodies against outer segment proteins (green). Rhodopsin clone 4D2 (**Da**), Red/Green opsin (**Db**), Wheat germ agglutinin (WGA, **Dc**). Nuclei are labelled with DAPI (4′,6-diamidino-2-phenylindole) in blue. Images are maximum intensity z-projections of 20 images taken at 0.7 µm intervals. OS: outer segment; IS: inner segment; ONL: outer nuclear layer, OPL: outer plexiform layer; INL: inner nuclear layer. Scale bars = 20 µm. Protein intensity profiles (green) along white lines shown at right. Magenta arrows: outer segment proteins mislocalized to inner segment or outer nuclear layer. Magenta asterisks mark outer segment proteins mislocalized to outer nuclear layer. n=4 animals examined by immunofluorescence microscopy. Raw data underling the graphs are available in S5 Data.

### Ift43 is required for assembly of full-length functioning cilia

To understand the importance of Ift43 in ciliary assembly, we generated fibroblasts from the *Ift43^tm1b/tm1b^* embryos and immortalized them with the Large T antigen (Figs 7-9). For comparison, we examined an *Ift140* control and homozygous mutant pair (Jonassen et al., 2012) alongside the *Ift43* cells. 50-80% of the cells of the controls for both *Ift140* and *Ift43* lines were ciliated with an average length of about 3 µm. Very few of the *Ift140* mutant cells ciliated but the cilia that assembled averaged 1.2 µm in length. The *Ift43* mutant cells were better ciliated with about 20% having cilia that averaged 1.4 µm in length (Fig 7E, 7F). As seen in the lymphatic cilia, the *Ift43* mutant cilia failed to label with Arl13b antibodies (S6 Fig), requiring the use of acetylated tubulin as the ciliary marker for most studies. To improve detection of cilia, acetylated tubulin staining was combined with either Cep164, which labels the distal appendages at the base of the cilium, or gamma tubulin, which labels the centrosome. The ciliary base marker used depended on the species and isotype of other antibodies used in an experiment.

**Figure 7.**
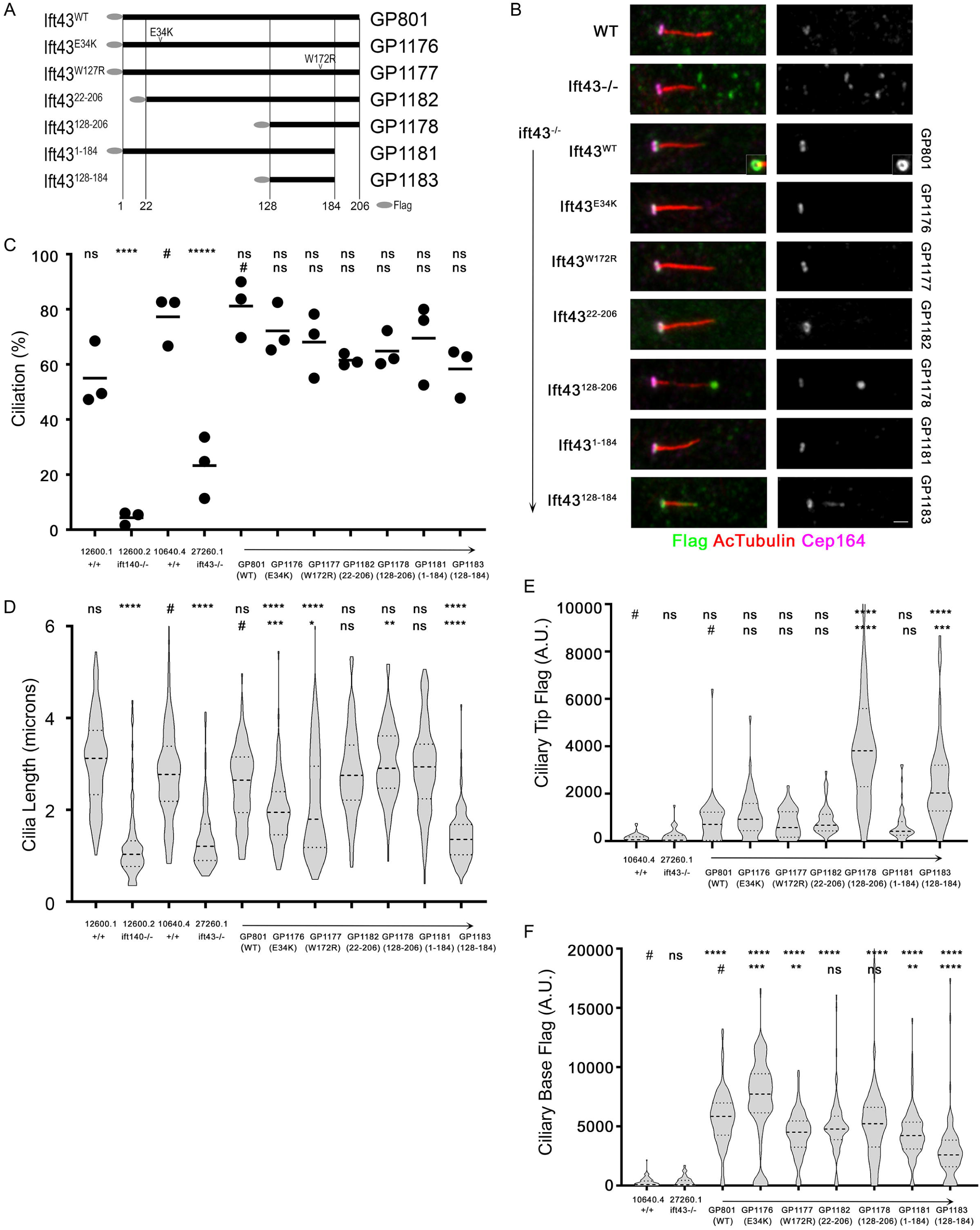
Ift43 is required for cilia formation. **(A)** Diagram of the Ift43 variants analyzed in this work. **(B)** Cilia stained for acetylated α-tubulin (AcTub, red), Cep164 (fuchsia), and Flag (green [left] or grey [right]) in wild type (10640.4), *Ift43* mutant (27260.1), and *Ift43* mutant fibroblasts rescued with wild type Ift43 tagged with Flag at the N-terminus (GP801) or the variants described in **A**. The inset in the wild-type rescue shows the base of a second cilium to illustrate that the Ift43 signal at the base co-localizes with Cep164 in a torus. Images are maximum projections of stacks taken at 0.14 µm intervals. Scale bar = 1 µm. **(C-D)** Percent ciliation (**C**) and cilia length (**D**) measured on serum starved MEFs. n>100 for each line and for ciliation, was repeated with three independently grown samples. ****p≤0.0001, ns not significant as compared to *Ift43* wild type (#, top row) or *Ift43* mutant rescued with wild-type Flag-Ift43 (#, bottom row) by one-way ANOVA with Tukey’s multiple comparisons post-hoc test. *Ift140* control and mutant are shown for comparison. Line in **C** represents mean, violin plots in **D** show median (bold dashed line) and quartiles (dashed lines). Note that some outliers are not plotted but all data was used for statistical analysis and is available in S5 Data. **(E-F)** Quantification of the Flag pools at the tip (**E**) and base (**F**) of cilia. The quantification tools of ImageJ were used to measure fluorescence intensity in an approximately 0.5-µm oval at the base or tip of each cilium. n>100 for each condition. Violin plots show median (bold dashed line) and quartiles (dashed lines). Note that some outliers are not plotted but all data was used for statistical analysis and is available in S5 Data. ****p≤0.0001, ***p≤0.001, **p≤0.01, ns not significant as compared to wild-type (#, top row) or *Ift43* mutant rescued with wild-type Flag-Ift43 (#, bottom row) by one-way ANOVA with Tukey’s multiple comparisons post-hoc test.

Both the cilia length and percent ciliation defects in the *Ift43* mutant cells were rescued by re-expression of wild-type *Ift43* tagged with Flag at the N-terminus (Fig 7A, 7E, 7F). When the rescue cells were stained for Flag, small amounts of signal could be observed along the shaft and at the tip, but the majority of the protein was localized at the base of the cilium where it colocalized with Cep164 in a torus (Fig 7B-7D). As described in the introduction, *Ift43* pathogenic variants are found in patients with isolated retinal degeneration (Ift43^E34K^) or skeletal dysplasias (Ift43^W179R^ [Ift43^W172R^ in mouse] and Ift43^M1X^). To determine how these variants affect ciliary function, we produced the equivalent mutations in the mouse protein (Fig 7A). The start codon variants were modeled by deleting the first 21 residues (Ift43^22-206^), which produces the protein expected if translation starts at the second methionine codon. This is not an exact mimic as the translational efficiency of message from this construct is likely better than the translational efficiency of message generated by the endogenous gene. In addition, we created a series of deletions based on the cryo structures of IFT-A (Hesketh et al., 2022; Jiang et al., 2023; Ma et al., 2023; Meleppattu et al., 2022). In these structures, only residues 128-184 are embedded into the particle. In the human structure the N-terminal 127 and C-terminal 22 residues are not observed, indicating that they are less ordered and may have functions other than holding the complex together. This fragment is contained within the portion of Ift43 that was sufficient to rescue ciliation in a *Chlamydomonas ift43* mutant (Zhu et al., 2017) and is conserved between species (S1 Data). Transfections of all of these constructs rescued the ciliation defect seen in the *Ift43* mutant cells, although the cilia on Ift43^E34K^ and Ift43^W172R^ were slightly shorter than controls (2.0 and 2.2 µm respectively) and the cells expressing Ift43^128-184^ had cilia about the same length (1.4 µm) as the *Ift43* mutant cells (Fig 7E, 7F). The ciliary localization of most variants was fairly similar to the controls with the protein found at the base and only a small amount along the shaft and at the tip. However, constructs missing the first 127 residues (Ift43^128-206^ and Ift43^128-184^) had significant tip accumulation (Fig 7B-7D).

The tip accumulation of Ift43 lacking the first 127 amino acids could indicate that this protein is not as effectively incorporated into retrograde particles or that the lack of this region generally disrupts IFT. To examine these possibilities, we quantified the effects of the mutant and deletion series on Ift88, an IFT-B protein, Ift140, an IFT-A protein, and Bbs5, a BBSome component (Fig 8). The loss of *Ift43* caused accumulation of Ift88 in the cilia (Fig 8A, B) and caused a redistribution of Ift140 from the basal body region to the ciliary tip (Fig D-F). It is difficult to observe Bbs5 in cilia unless retrieval is disrupted by an *Ift27* mutation (Eguether et al., 2014). Thus, as a control, we included an *Ift27* mutant MEF in the experiment. As expected, nearly all of the *Ift27* mutant cells had robust Bbs5 label in cilia but the control cilia were not labeled. While less robust than what is seen by the loss of *Ift27*, the loss of *Ift43* significantly increased the number of cilia where Bbs5 was detected. Typically, the label was in a punctum at the tip but also could be found along the cilia (Fig 8H, I). The Ift88 ciliary accumulation phenotype is also seen in the *ift140* mutant cells (Fig 8C), but Bbs5 was not enriched in cilia of *Ift140* mutant cells (Fig 8J). As expected, Ift140 was not observed in *Ift140* mutant cells (Fig 8G). The Ift88 ciliary tip accumulation was rescued by all of the variants except for Ift43^128-184^, which maintained a slight enrichment at the tip. The Ift140 tip accumulation and Bbs5 ciliary accumulation phenotypes were rescued by all constructs except those missing the N-terminal 127 residues (Ift43^128-206^ and Ift43^128-184^). These are the two constructs that showed enrichment in cilia (Fig 7D) suggesting the IFT particles assembled without the N-terminus of Ift43 are less efficiently trafficked out of cilia. This finding also suggests that Ift43 plays a role in retrograde movements of the BBSome, a function not previously known.

**Figure 8.**
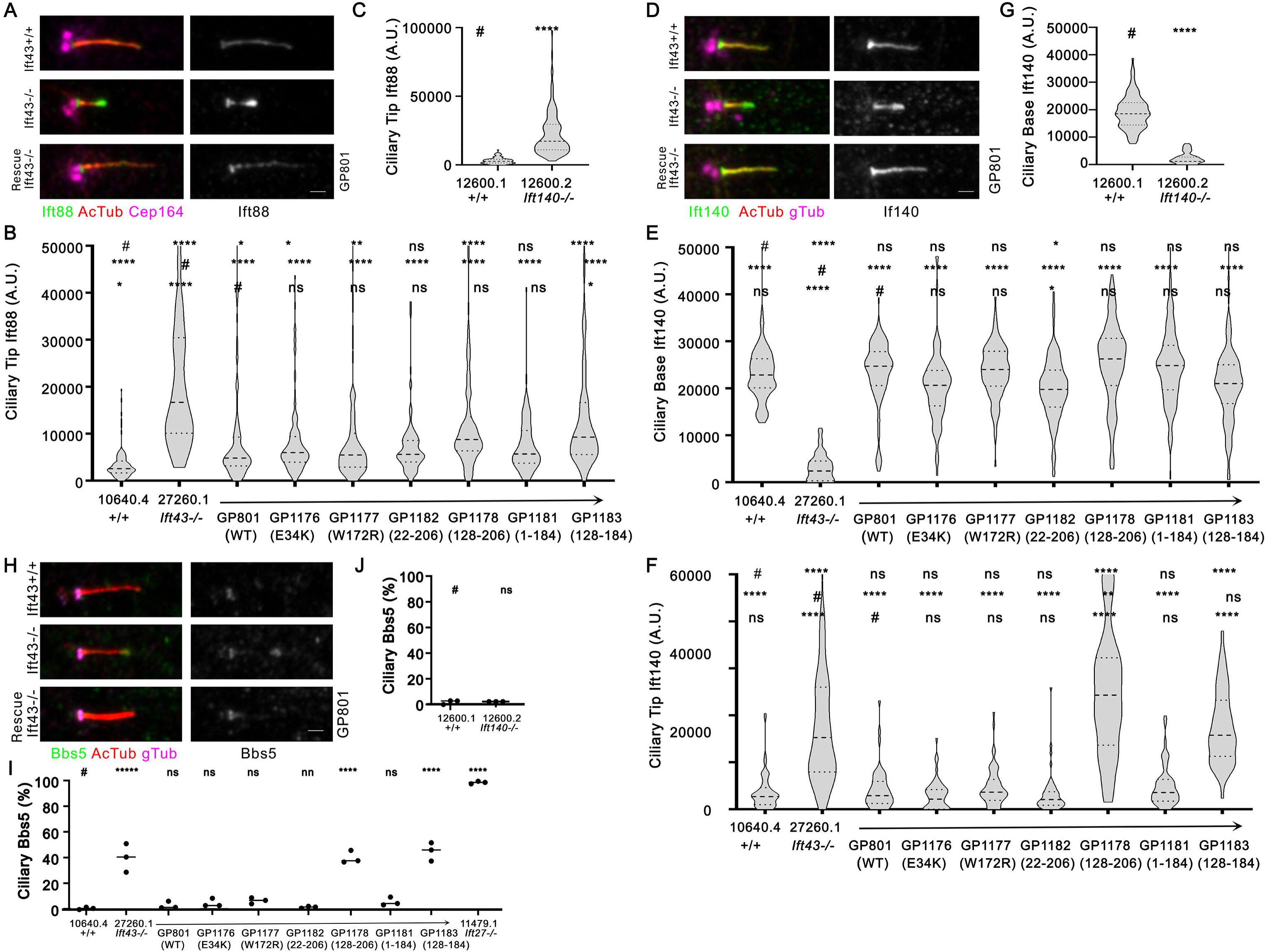
Ift88, Ift140, and Bbs5 accumulate in *Ift43* mutant cilia. **(A)** Cilia stained for acetylated α-tubulin (AcTub, red), Cep164 (fuchsia), and Ift88 (green [left] or grey [right]) in wild type (10640.4), *Ift43* mutant (27260.1), and *Ift43* mutant fibroblasts rescued with wild type Ift43 tagged with Flag at the N-terminus (GP801). Images are maximum projections of stacks taken at 0.14 µm intervals. Scale bar = 1 µm. **(B)** Quantification of the Ift88 pool at the ciliary tip of *Ift43* MEFs. The quantification tools of ImageJ were used to measure fluorescence intensity in an approximately 0.5-µm oval at the tip of each cilium. n>100 for each condition. Violin plots show median (bold dashed line) and quartiles (dashed lines). Note that some outliers are not plotted but all data was used for statistical analysis and is available in S5 Data.****p≤0.0001, **p≤0.01, *p≤0.05, ns not significant as compared to wild-type (#, top row), *ift43* mutant (#, middle row) or wild-type rescue (#, bottom row) by one-way ANOVA with Tukey’s multiple comparisons post-hoc test. **(C)** Quantification of the Ift88 pool at the ciliary tip of *Ift140* MEFs. Parameters are the same as described in (**B)**. Raw data underling the graph is available in S5 Data. **(D)** Cilia stained for acetylated α-tubulin (AcTub, red), Cep164 (fuchsia), and Ift140 (green [left] or grey [right]) in wild type (10640.4), *Ift43* mutant (27260.1), and *Ift43* mutant fibroblasts rescued with wild type Ift43 tagged with Flag at the N-terminus (GP801). Images are maximum projections of stacks taken at 0.14 µm intervals. Scale bar = 1 µm. **(E-F**) Quantification of the Ift140 pool at the ciliary base (**E**) or tip (**F**) of *Ift43* MEFs. The quantification tools of ImageJ were used to measure fluorescence intensity in an approximately 0.5-µm oval at the base or tip of each cilium. n>100 for each condition. Violin plots show median (bold dashed line) and quartiles (dashed lines). Note that some outliers are not plotted but all data was used for statistical analysis and is available in S5 Data. ****p≤0.0001, **p≤0.01, *p≤0.05, ns not significant as compared to wild-type (#, top row), *ift43* mutant (#, middle row) or wild-type rescue (#, bottom row) by one-way ANOVA with Tukey’s multiple comparisons post-hoc test. **(G)** Quantification of the Ift140 pool at the ciliary base of *Ift140* MEFs. Parameters are the same as described in (**E)**. Raw data underling the graph is available in S5 Data. **(H)** Cilia stained for acetylated α-tubulin (AcTub, red), γ-tubulin (gTub, fuchsia), and Bbs5 (green [left] or grey [right]) in wild type (10640.4), *Ift43* mutant (27260.1), and *Ift43* mutant fibroblasts rescued with wild type Ift43 tagged with Flag at the N-terminus (GP801). Images are maximum projections of stacks taken at 0.14 µm intervals. Scale bar = 1 µm. **(I)** Percent of *Ift43* cilia with a detectable Bbs5 puncta in cilia. n = 3 independent repeats with >100 cilia counted per condition. Line represents mean. ****p≤0.0001, ns not significant as compared to *Ift43* wild type (#) by one-way ANOVA with Tukey’s multiple comparisons post-hoc test. Raw data underling the graphs are available in S5 Data. **(J)** Percent of *Ift140* cilia with a detectable Bbs5 puncta in cilia. Parameters are the same as described in (I). Raw data underling the graph is available in S5 Data.

### Ift43 is required for efficient Hedgehog signaling

Induction of Hedgehog signaling promotes the production of Gli1 protein and reduction in the processing of Gli3 full length (Gli3^FL^) to the repressor form (Gli3^R^) (Wang et al., 2000). Gli2 is also processed similar to Gli3 into repressor forms but to a lesser extent (Bhatia et al., 2006; Pan et al., 2006). To determine the importance of Ift43 to Hedgehog signaling, we used western blotting to examine the levels and processing of Gli1, Gli2, and Gli3 before and after SAG stimulation. As expected, control cells show low Gli1 at the basal state with an increased level after agonist treatment (Fig 9A). In controls, Gli2^FL^ levels are approximately equal at basal and induced conditions and the repressor form is barely detectable at the basal state (Fig 9B). In controls at the basal state, Gli3^FL^ and Gli3^R^ are observed at approximately equal levels. With activation, Gli3^R^ is mostly gone and Gli3^FL^ is slightly reduced (Fig 9C). SAG treatment of the *Ift43* mutant cells fails to increase Gli1 protein levels (Fig 9A), fails to stop the processing of Gli3^FL^ to Gli3^R^ (Fig 9C), and fails to stop the minimal processing of Gli2^FL^ to Gli2^R^ normally observed (Fig 9B) The Gli1 and Gli3 phenotypes were rescued by the expression of wild-type Flag-tagged protein (GP801) and all of the patient and deletion variants. Curiously, the overexpression of wild-type Flag-tagged Ift43 and all the variants increased the processing of Gli2^FL^ into Gli2^R^ at the basal state (Fig 9B) suggesting that Ift43 is limiting for the cleavage of Gli2.

**Figure 9.**
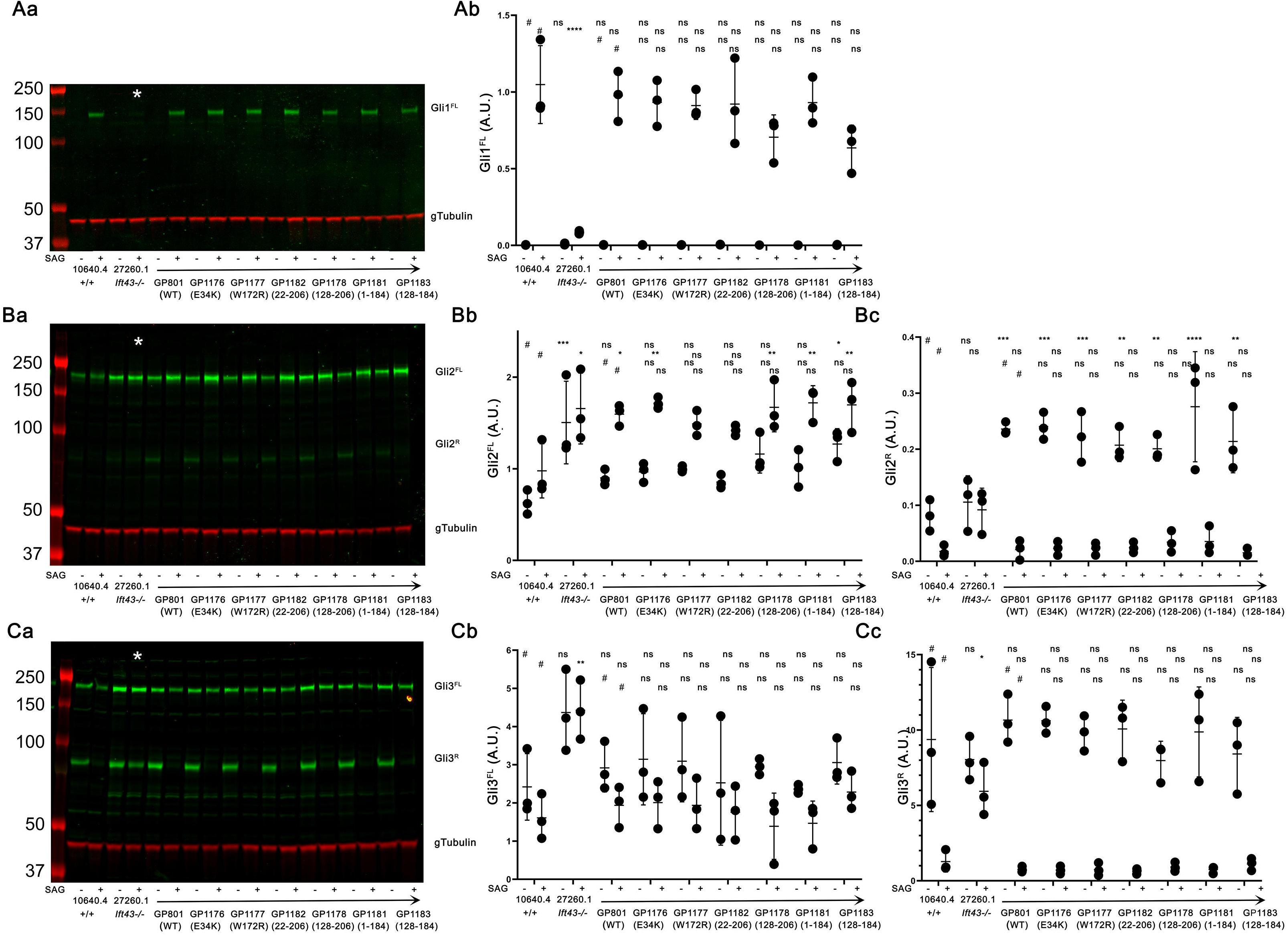
Ift43 is required for Hedgehog Signaling. **(Aa)** Western blot showing Gli1 protein (green) in unstimulated (-) or SAG stimulated (+) wild type (10640.4), *Ift43* mutant (27260.1), and *Ift43* mutant fibroblasts transfected with wild-type *Ift43* or variants described in Fig 7A. γ-tubulin (red) was used as a loading control. * marks the lane containing *Ift43* mutant stimulated with SAG. **(Ab)** Quantification of the Gli1 band from three biological replicates. Bars are mean and standard deviation. ****p≤0.0001, ns: not significant by two-way ANOVA with Tukey’s multiple comparisons test. **(Ba)** Western blot showing Gli2 protein (green) in unstimulated (-) or SAG stimulated (+) wild type (10640.4), *Ift43* mutant (27260.1), and *Ift43* mutant fibroblasts transfected with wild-type *Ift43* or variants described in Fig 7A. γ-tubulin (red) was used as a loading control. * marks the lane containing *Ift43* mutant stimulated with SAG. **(Bb**) Quantification of the Gli2^FL^ uncleaved band from three biological replicates. Bars are mean and standard deviation. ***p≤0.001, **p≤0.01, *p≤0.05, ns: not significant by two-way ANOVA with Tukey’s multiple comparisons test. Raw data underling the graphs are available in S5 Data. **(Bc)** Quantification of the Gli2^R^ cleaved band from three biological replicates. Bars are mean and standard deviation. ****p≤0.0001, ***p≤0.001, **p≤0.01, ns: not significant by two-way ANOVA with Tukey’s multiple comparisons test. **(Ca)** Western blot showing Gli3 protein (green) in unstimulated (-) or SAG stimulated (+) wild type (10640.4), *Ift43* mutant (27260.1), and *Ift43* mutant fibroblasts transfected with wild-type *Ift43* or variants described in Fig 7A. γ-tubulin (red) was used as a loading control. * marks the lane containing *Ift43* mutant stimulated with SAG. **(Cb)** Quantification of the Gli3^FL^ uncleaved band from three biological replicates. Bars are mean and standard deviation. **p≤0.01, ns: not significant by two-way ANOVA with Tukey’s multiple comparisons test. **(Cc**) Quantification of the Gli3^R^ cleaved band from three biological replicates. Bars are mean and standard deviation. *p≤0.05, ns: not significant by two-way ANOVA with Tukey’s multiple comparisons test. Uncropped western blots are available in S5 Data and raw data underling the graphs are available in S5 Data.

To understand the step at which the pathway is blocked in the *Ift43* mutant and rescue cells, we quantified ciliary Smo, Gli2, and Gli3 levels at basal and SAG-induced states. The ciliary level of Smo at the basal state was not different in the *Ift43* mutant cells compared to controls but the increase in ciliary level after SAG stimulation was muted (Figs 10A, 10B). *Ift140* mutant cells also failed to increase ciliary Smo upon SAG treatment (Fig 10C). The failure to enrich Smo in cilia after SAG treatment phenotype could be rescued by expression of wild-type Flag-tagged protein. The pathogenic and deletion variants showed a mix of decreased (Ift43^E34K^, Ift43^W172R^, Ift43^22-206^) and increased (Ift43^128-206^, Ift43^1-184^, Ift43^128-184^) enrichment after SAG stimulation (Fig 10B) although they all generally elevated Smo levels with SAG treatment.

**Figure 10.**
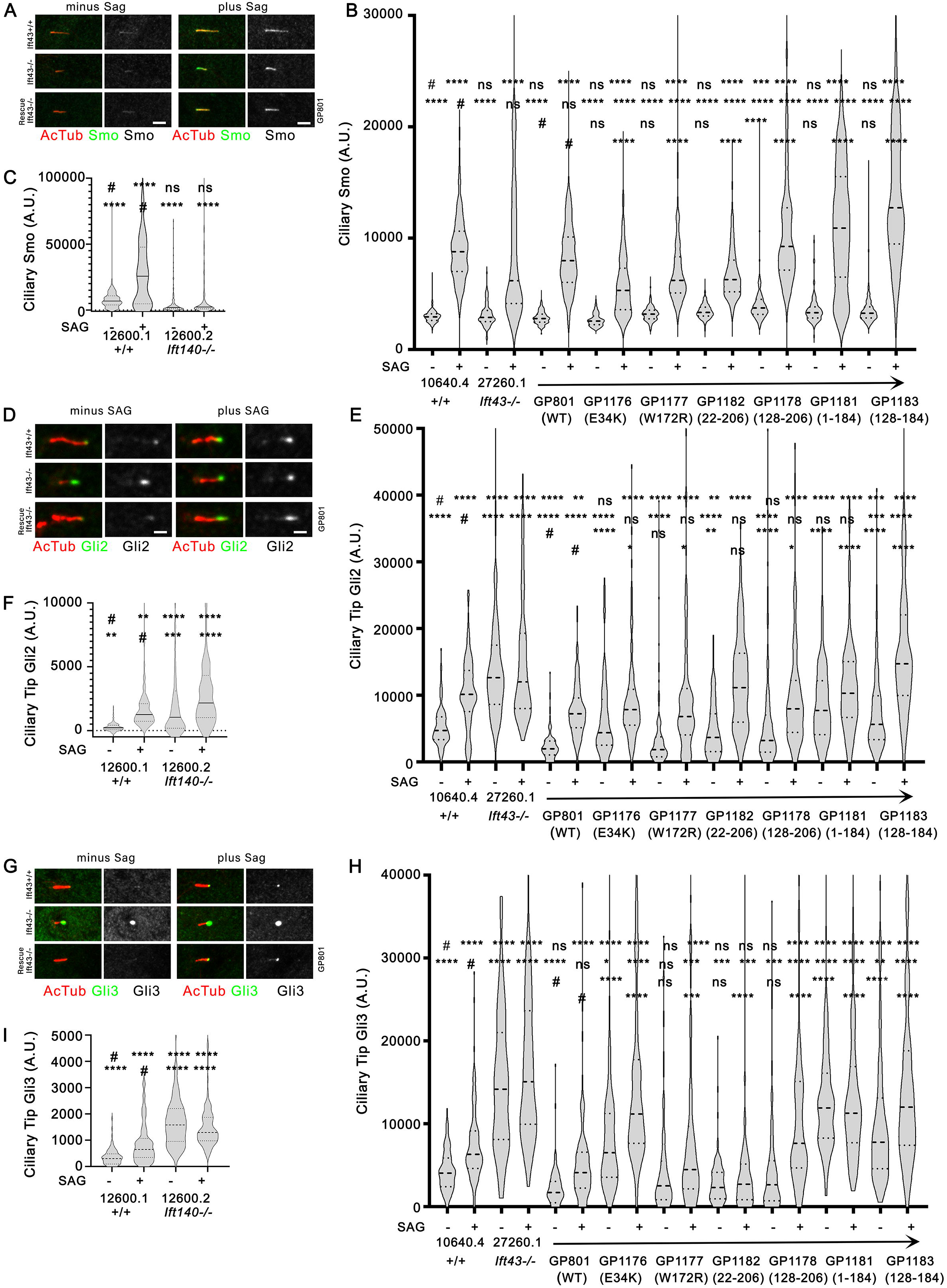
Ift43 mutants accumulate Gli2 and Gli3 inappropriately at the ciliary tip. **(A)** Cilia stained for acetylated α-tubulin (AcTub, red) and Smo (green [left], grey [right]) in unstimulated (minus SAG) or SAG stimulated (plus SAG) wild type (10640.4), *Ift43* mutant (27260.1), and *Ift43* mutant fibroblasts rescued with wild type Ift43 tagged with Flag at the N-terminus (GP801). Images are maximum projections of stacks taken at 0.19 µm intervals. Scale bars = 1 µm. **(B)** Quantification of ciliary Smo in unstimulated (-) or SAG stimulated (+) wild type (10640.4), *Ift43* mutant (27260.1), and *Ift43* mutant fibroblasts transfected with wild-type *Ift43* or variants described in Fig 7A. The quantification tools of CiliaQ were used to measure fluorescence intensity. n>100 for each condition. Violin plots show median (bold dashed line) and quartiles (dashed lines). Note that some outliers are not plotted but all data was used for statistical analysis and is available in S5 Data. ****p≤0.0001, ns not significant as compared to wild type without and with SAG (#, top two rows) or *ift43* mutant rescued with wild type Flag-Ift43 without and with SAG (# bottom two rows) by two-way ANOVA with Tukey’s multiple comparisons post-hoc test. **(C)** Quantification of ciliary Smo in *Ift140* MEFs. Parameters are the same as described in **(B)**. Raw data underling the graph is available in S5 Data. **(D)** Cilia stained for acetylated α-tubulin (AcTub, red) and Gli2 (green [left], grey [right]) in unstimulated (minus SAG) or SAG stimulated (plus SAG) wild type (10640.4), *Ift43* mutant (27260.1), and *Ift43* mutant fibroblasts rescued with wild type Ift43 tagged with Flag at the N-terminus (GP801). Images are maximum projections of stacks taken at 0.19 µm intervals. Scale bars = 1 µm. **(E)** Ciliary tip levels of Gli2 in unstimulated (-) or SAG stimulated (+) wild type (10640.4), *Ift43* mutant (27260.1), and *Ift43* mutant fibroblasts transfected with wild-type *Ift43* or variants described in Fig 7A. Violin plots show median (bold dashed line) and quartiles (dashed lines). Note that some outliers are not plotted but all data was used for statistical analysis and is available in S5 Data. **** p≤0.0001, *** p≤0.001, ** p≤0.01, * p≤0.05, ns not significant by two-way ANOVA with Tukey’s multiple comparison test as compared to wild type without and with SAG (#, top two rows) or *ift43* mutant rescued with wild type Flag-Ift43 without and with SAG (# bottom two rows). n>100 cilia per condition. **(F)** Ciliary tip levels of Gli2 in wild type (12600.1) and *Ift140* mutant (12600.2) MEFs. Parameters are the same as described in **(E)**. Raw data underling the graph is available in S5 Data. **(G)** Ciliary tip levels of Gli3 in unstimulated (-) or SAG stimulated (+) wild type (10640.4), *Ift43* mutant (27260.1), and *Ift43* mutant fibroblasts transfected with wild-type *Ift43* or variants described in Fig 7A. Violin plots show median (bold dashed line) and quartiles (dashed lines). Note that some outliers are not plotted but all data was used for statistical analysis and is available in S5 Data. **** p≤0.0001, *** p≤0.001, ** p≤0.01, * p≤0.5, ns not significant by two-way ANOVA with Tukey’s multiple comparison test as compared to wild type without and with SAG (#, top two rows) or *ift43* mutant rescued with wild type Flag-Ift43 without and with SAG (# bottom two rows). n>100 cilia per condition.

At the basal state, Gli2 is normally found at low levels at the tip of cilia and becomes enriched at this site upon pathway activation. In *Ift43* mutant cells, Gli2 is significantly enriched at the ciliary tip at the basal state and is not greatly enriched by SAG treatment (Figs 10D, 10E). This basal-state enrichment is rescued by re expression of wild type. The patient variants and deletions showed intermediate phenotypes with more Gli2 at the ciliary tip than controls but not as much as the *Ift43* mutant (Fig 10E). *Ift140* mutants also show basal-state Gli2 enrichment at the ciliary tip (Fig 10F). These results suggest that Ift43 and Ift140, probably acting together as part of IFT-A, function in removing Gli2 from cilia at the basal state.

Similar to Gli2, Gli3 is also enriched at the ciliary tip after pathway activation (Fig 10G). The Ift43 mutant cilia show increased basal levels of Gli3 at the tip and this phenotype is rescued by re expression of wild type Flag-Ift43. Similar to what was observed with Gli2, the patient and deletion mutants showed intermediate phenotypes with the patient variant Ift43^E34K^ and the C-terminal deletions of 22 residues (Ift43^1-184^ and Ift43^128-184^) causing the largest increase in ciliary tip Gli3 (Fig 10G). The accumulation of Gli2 and Gli3 at the ciliary tip under basal conditions suggests that the ciliary level is controlled by regulated removal rather than regulated delivery.

## Discussion

Ift43, the least conserved subunit of IFT-A is critical to the formation and function of cilia. In mice, we found that a null allele leads to mid-to-late gestational lethality with about 50% of homozygotes dead by E15.5. This is slightly less severe than we observed with an *Ift140* null allele where more than 50% died by E13.5. This less severe phenotype is consistent with the observation that very few *Ift140* mutant fibroblasts formed cilia, while about 20% of *Ift43* cells formed cilia. The *Ift43* mutant embryos showed severe craniofacial abnormalities with a large gaping mouth and a small tongue. Brain malformations including exencephaly and holoprosencephaly were observed. Eye development appeared to be arrested at an early state with most animals showing no eye development and only one animal showing evidence of an eye cup. In addition to a role in early eye development, Ift43 is crucial for photoreceptor outer segment maintenance as deletion of an *Ift43* floxed allele with photoreceptor rod-specific Cre led to degeneration of the outer segment and accumulation of vesicular material in the inner segment. The mutants showed extreme hydrops where fluid accumulated under the skin. This type of fluid accumulation is typically due to defective lymphatic function, which was consistent with our finding that lymphatic vessels were abnormally large. The mutants showed omphalocele and cordis ectopia due to a body wall closure defect allowing the heart, liver, and other organs to be exposed. The body wall defect is accompanied by a diaphragmatic hernia allowing the thoracic and abdominal organs to intermingle with the liver projecting up and around the lungs and heart. The lungs were strongly affected with only a single hypoplastic central lung observed. Cardiac phenotypes included a common atrium where the septa separating the left and right atrium is missing preventing the separation of systemic and pulmonary blood supplies, persistent truncus arteriosus where the aorta and pulmonary artery fail to separate into two separate vessels, dextroversion where the heart is rotated clockwise placing the left ventricle in front of the right ventricle (Grant, 1958) and superior inferior ventricles where the right ventricle is located above the left.

In vertebrates the hedgehog pathway is organized around the primary cilium and mutations affecting IFT components typically block the pathway but the role of the IFT proteins in hedgehog signaling is complex. They are needed for the assembly of cilia where the signaling occurs, but hypomorphic *Ift* mutations that can partially form cilia have defective signaling (Ocbina et al., 2011) suggesting that IFT proteins have signaling roles in addition to their function in assembly. We previously showed that the IFT-B proteins Ift25 and Ift27 are not required for ciliary assembly but cells that built cilia lacking these proteins are defective in hedgehog signaling similar to cells lacking cilia (Eguether et al., 2018; Eguether et al., 2014; Keady et al., 2012). The defect in hedgehog signaling reflects the need of these two IFT subunits to remove Smo from cilia at the basal state and remove Ptch1 from cilia after pathway induction. In these mutants the delivery of Gli2 to the cilium in response to SAG treatment was normal, but it failed to accumulate at the ciliary tip. We originally focused on Ift25 and Ift27 as they are missing from *Drosophila* and *Caenorhabditis*. Since these organisms that can form cilia without these two proteins, we reasoned that they may be involved in a ciliary function not needed in these organisms. In *Drosophila*, hedgehog signaling is cilia independent while *Caenorhabditis* does not have a classic hedgehog pathway, consistent with a role for Ift25 and Ift27 in hedgehog signaling. We noted that Ift43 was often lost from organisms that lost Ift25 and Ift27 suggesting that this IFT-A protein might likewise be more important for ciliary function than for ciliary assembly. In contrast to *Ift25* and *Ift27* mutants that form normal numbers of full length or longer cilia, *Ift43* mutants have a reduced number of shorter cilia. This is likely due to the requirement of Ift43 to bridge Ift121 to Ift139 within IFT-A (Hesketh et al., 2022; Jiang et al., 2023; Ma et al., 2023; Meleppattu et al., 2022). Mutants that are missing Ift43 accumulate Ift88, Ift140, and Bbs5 consistent with a general disruption of IFT. The re-expression of the structural portion of Ift43 is sufficient to rescue the decrease in ciliation seen in the mutant line. However, this fragment is not sufficient to fully rescue the length phenotype, suggesting that the disordered N- or C-terminal ends have additional functions.

Similar to what we observed in *Ift25* and *Ift27* mutants, we find that *Ift43* mutants fail to activate Hedgehog signaling in response to SAG. Loss of *Ift25* or *Ift27* caused an inappropriate accumulation of Smo in cilia at the basal state and a failure to concentrate Gli2 at the ciliary tip after activation. We do not observe Smo accumulation with the loss of *Ift43*. Instead, we found that the basal levels of Gli2 and Gli3 at the ciliary tip are higher than normal in the *Ift43* mutant. This finding suggests that the level of Gli2 and Gli3 at the ciliary tip is regulated by removal driven by Ift43. In this role, Ift43 is probably acting as part of IFT-A as *Ift140* mutants also show increased ciliary Gli2 at the basal state. Gli2 is actively transported by IFT (Ku et al., 2025) so a direct role for Ift43 and IFT-A in transporting Gli2 is possible. *Ift43* mutants fail to elevate Gli1 protein in response to SAG activation despite being able to enrich Smo. Normally, activation by SAG blunts the production of Gli3^R^ and the promotion of active forms of Gli2 and Gli3. Cells lacking *Ift43* continue to produce the cleaved Gli3^R^ suggesting that the continued production of the repressor may be the reason for the lack of Gli1 upregulation with SAG treatment. Normally Gli2 is maintained in an inactive state by sequestration in complex with Sufu but a small amount is processed into the cleaved repressor form. This cleavage continues after SAG treatment in *Ift43* mutants, which may provide more repression of Hedgehog targets. Interestingly, the overexpression of Ift43 increases the production of Gli2^R^ suggesting that Ift43 is limiting for the processing of Gli2. The processing of the Gli transcription factors to repressor forms is thought to be the result of a cascade initiated by PKA phosphorylation in response to high cAMP levels, followed with ubiquitylation by β-TRCP (Bhatia et al., 2006; Pan et al., 2006) and ultimately cleavage by the proteosome. When the pathway is activated, cAMP levels drop reducing PKA phosphorylation of Gli and its subsequent cleavage into the repressor form. In addition, alternative sites become phosphorylated, which elevates transcription-promoting activity (Niewiadomski et al., 2014). The continued cleavage suggests that PKA activity is maintained likely due to a failure to suppress cAMP levels with SAG treatment.

Our finding that Gli2 and Gli3 repressor forms continue to be produced after pathway activation suggests that the *Ift43* mutation causes reduced signaling through the pathway. This phenotype may be expected to mimic mutations which reduce the production of the Hedgehog ligands, the loss of Smo, or mutations that express only the repressor forms of Gli2 and Gli3. While the loss of SHH or Smo produces phenotypes like holoprosencephaly and oligodactyly and syndactyly (Chiang et al., 1996; Zhang et al., 2001), the overall phenotype is more extreme than we observed with Ift43 indicating that the loss of Ift43 is not completely blocking the pathway. Pallister-Hall syndrome is caused by variants in Gli3 that truncate the protein after the zinc-finger DNA binding domain and are thought to produce Gli3 with only repressor activity (Demurger et al., 2015). This disease is typically *de novo* due to new mutations on one chromosome or is inherited dominantly. The patients present with a variety of limb abnormalities including mesoaxial polydactyly, postaxial polydactyly, bifid epiglottis, and hypothalamic hamartomas along with other phenotypes that can include imperforate anus, renal abnormalities including cystic disease and hypoplasia, ectopic ureteral implantation, genitourinary anomalies, pulmonary segmentation anomalies including bilateral bilobed lungs, and short limbs (Demurger et al., 2015). Two mouse models of Pallister-Hall syndrome have been produced. The first called *Gli3*^Δ*699*^ produced a protein truncated after the zinc finger consisting of the first 699 residues of Gli3 with 21 residues of thymidine kinase fused to the C-terminus (Bose et al., 2002). The second allele called *Gli3*^Δ*701C*^ also produced a protein that after Cre excision is truncated after the zinc finger and consists of the first 701 Gli3 residues (Cao et al., 2013). While *Gli3*^Δ*699*^ was adult viable, *Gli3*^Δ*701C*^ appeared to be embryonic lethal as no offspring were obtained when the gene was deleted with a developmentally active, ubiquitously expressed Cre driver even in the heterozygous state. Homozygotes of *Gli3*^Δ*699*^ or heterozygotes of *Gli3*^Δ*701C*^ deleted in limb lineages showed digit phenotypes like what we observed in Ift43 mutants consistent with our hypothesis that the limb phenotype in *Ift43* mutants is driven by the increased production of Gli3^R^ and possibly Gli2^R^.

In summary, our work shows that while Ift43 is not completely required for ciliary assembly, the cilia assembled on mutant cells are fewer in number, shorter in length, and fail to induce the Hedgehog pathway in response to agonist. The failure to induce the pathway is likely due to an inability to suppress the conversion of Gli2 and Gli3 into cleaved repressor forms. Unexpectedly, the over expression of Ift43 caused increased processing of Gli2 into the repressor form, indicating that Ift43 is limiting for this process. In addition, the loss of Ift43 caused elevated levels of Gli2 and Gli3 at the ciliary tip, suggesting that like Ptch1 and Smo, the ciliary levels of the Gli transcription factors are controlled by regulated removal from cilia.

## Experimental Procedures

### Mice

Mouse work was carried out at the University of Massachusetts Chan Medical School with IACUC approval. Strain 037736-UCD, C57BL/6N-^Atm1Brd^/a *Ift43^tm1a(EUCOMM)Hmgu^/BcmMmucd* was obtained from the Mouse Biology Program (University of California, Davis). *Ift43^tm1a^* mice were converted to *Ift43^tm1b^* by crossing to germline Zp3-Cre (003651, C57BL/6-Tg(Zp3-cre)93Knw/J, Jackson Laboratories, Bar Harbor ME USA) and to the *Ift43^tm1c^* (flox) alleles by crossing to FlpO (B6.Cg-Tg(Pgk1-flpo)10Sykr/J, Jackson Laboratories, 003651) The lines were maintained by recurrent mating to wild type C57Bl/6J (000664, Jackson Laboratories). Genotyping details are provided in S2 Data.

### Optical Projection Tomographic Imaging

Embryos were fixed in 4% paraformaldehyde for at least 24 hr, then briefly washed with phosphate-buffered saline and embedded in 1.1% w/v low melting point agarose in MilliQ water. Agarose-embedded embryos were then mounted to magnetic stubs using Loctite 454 surface-insensitive instant adhesive glue, dehydrated in 100% methanol for three days, changing to fresh methanol each day. Following dehydration, embedded embryos were cleared in BABB (2:1, benzyl alcohol:benzyl benzoate), changing the BABB each day until the embryos were clear. Cleared, embedded embryos were then imaged using a Bioptonics 3001M Optical Projection Tomograph under UV light with a GFP1 filter (425nm/40nm, 475nmLP). Raw scan images were then reconstructed into multiplanar slice data using NRecon software (V1.7.4.6) operated through the NRecon GPU Server and then imported into Drishti Volume Exploration software (v3.2) for visual assessment.

### Cell Culture

Mutant MEFs were isolated from E13.5 mutant embryos and immortalized with the large T antigen from SV40 virus. MEFs and their derivatives were grown at 37°C in 5% CO_2_ in Dulbecco’s modified Eagle’s medium (11995-065, Gibco ThermoFisher, Waltham MA USA) with 5% FBS and 1% Penicillin-Streptomycin (25200-056, Gibco ThermoFisher). All DNA constructs were transfected into the cells via lentiviral infection followed by drug selection to create stable cell lines.

Cells for immunofluorescence microscopy were grown on acid-washed glass coverslips. The cells were fixed for 10 min in −20° C methanol or for 15 min in 2% paraformaldehyde, 0.05 M Pipes, 0.025 M Hepes, 0.01 M EGTA, 0.01 M MgCl2 (pH 7.2) followed by a two-min extraction with 0.1% Triton X-100 in the same solution. For some antibodies, an antigen retrieval step of 0.05% SDS in PBS for 5 minutes included at this point. After two brief washes in TBST (0.01 M Tris, pH 7.5, 0.166 M NaCl, 0.05% Tween 20), the cells were blocked with 1% bovine serum albumin (BSA, Sigma-Aldrich, St. Louis MO USA) in TBST for 1 hr and then incubated with the primary antibodies either overnight at 4°C or for 2 hr at room temperature. The cells were then washed 4 times with 1% BSA/TBST over ∼30 min. The cells were then incubated with 1:2000 dilutions of Alexa fluor-conjugated secondary antibodies (S2 Table) for 1 hr and washed 4 times with 1% BSA/TBST over ∼30 min followed by a brief wash with TBST. The cells were then mounted with ProLong Antifade with 4′,6-diamidino-2-phenylindole (DAPI, P36935, Invitrogen ThermoFisher, Waltham MA USA). Cells were imaged on an LSM 910+ Airyscan confocal microscope (Zeiss, Oberkochen Germany) with a 63× or 40x objectives. If comparisons are to be made between images, the photos were taken with identical conditions and manipulated equally.

For SAG experiments, cells were plated at near confluent densities and serum starved (same culture medium described above but with 0.25% FBS) for 48 hrs prior to treatment to allow ciliation. SAG (566660, Sigma-Aldrich, St. Louis MO USA) was used at 400 nM.

Cells were confirmed to be of mouse origin and monitored for mycoplasma contamination by PCR (Tang et al., 2000) and DAPI staining.

### Quantification of Ciliary Parameters

To measure the accumulation of Smo and Arl13b in cilia, we developed a workflow around the CiliaQ ImageJ plugin (Hansen et al., 2021). Cilia were first identified using the ilastik random forest image segmentation tool (Berg et al., 2019) and the cilia masks were added to the image stacks to mimic the output of CiliaQ preparator using ImageJ macros (S7 Data).These images with the embedded masks were subjected to the quantification tools of CiliaQ (module 3). Contamination in the data from midbodies and cytoplasmic microtubule bundles were removed by a blind approach. To achieve this, data from the control and experimental conditions were put through a custom ImageJ plugin that collected all of the regions of interest (ROI) in the dataset and presented them as random grids allowing the investigator to mark and remove non-ciliary contamination while being blinded to the source of the ROI (Anuszczyk et al., 2025) (code is available at https://github.com/pazourg/DeleteROI).

Ciliary tip or base Gli2, Gli3, Ift88, and Ift140 were determined using the quantification tools of ImageJ in a non-blinded approach. To do this, image stacks were converted to maximum projection single planes, an ROI was defined at the tip or base of the cilium, and the average intensity of the ROI and its area determined. The local background was measured next to the cilium tip. signal intensity = (area of ROI)*(average intensity of ROI - average intensity of local background).

### Lymphatic Analysis

E15.5 embryos were fixed using 4% paraformaldehyde. Dorsal skin was blocked and permeabilized overnight at RT with shaking in PBS^++^ (5.2% BSA, 0.3% Triton X-100, and 0.2% sodium azide in PBS). Primary and secondary antibodies (S3 Table) were applied in PBS^++^ overnight with shaking. Following each round of antibodies three one-hr washes of PBS^+^ (0.2% BSA, 0.3% Triton X-100, and 0.2% sodium azide) were carried out at room temperature with shaking. Samples were mounted as whole mounts in Fluoromount-G without DAPI (Southern Biotech, Birmingham AL USA, 0100-01) for imaging.

Images were captured by laser scanning confocal microscopy using a FLUOVEW FV1200 scanning confocal microscope (Olympus, Tokyo Japan) interfaced with an IX81 microscope (Olympus, Tokyo Japan) using 488, 568, and 647 laser lines. Z-step size was 0.5 or 1 μm as indicated in figure legends.

Lymphatic vessel density was quantified using the freehand selection tool in FIJI (NIH ImageJ). Maximum intensity projections were used to outline the vessel area. The total vessel area for each field of view was then divided by the total field of view area to achieve a percent vessel density.

Lymphatic endothelial cell proliferation was quantified using a FIJI and Cell Profiler. Merged 20X images acquired through confocal imaging were processed slice by slice to determine colocalization between Prox1 and phospho-histone H3. The multipoint selection tool was used to mark this colocalization. A Cell Profiler pipeline was configured to count total Prox1 nuclei using maximum intensity projections of the Prox1 channel. The pipeline was configured to identify primary objects, edit objects manually, save images, and export data to a spreadsheet. The diameter and intensity threshold parameters for identifying primary objects were optimized for each image. Pprox1 signals touching the border of the image were not counted. The number of proliferating nuclei based on the colocalization of PROX1 and phospho-histone H3 was then divided by the total number of Prox1 nuclei to achieve a percent proliferation of lymphatic endothelial cells per field of view.

E15.5 vessel diameter quantification was performed using FIJI. The grid tool was used to overlay a grid over each merged 10X maximum intensity projections (Grid type: Lines, Area per point: 19975.16 microns^2^). The segmented line tool was used to measure the diameter of each vessel that perpendicularly crossed the grid lines within 10° tolerance. Every diameter within the tolerance was measured.

### Retina Analysis

Electroretinogram (ERG) recordings were conducted using a Celeris system (Diagnosys, Lowell MA, USA) to evaluate retinal responses under combined dark- and light-adapted conditions, including a flicker response. After a 12 hr dark adaptation period, mice were anesthetized via an intraperitoneal injection of a ketamine-xylazine mixture (100 mg/kg of body weight and 10 mg/kg of body weight, respectively). Pupil dilation was achieved by applying one drop each of phenylephrine (2.5%) and tropicamide (1%) at ∼10 min before recording. Throughout the ERG procedure, animals were maintained on a warming plate to sustain a body temperature of 37°C. ERG electrodes were directly positioned on the eyes and a flicker ERG frequency series was performed with intensities of 0.01 cd·s/m^2^, 0.1 cd·s/m^2^, and 1 cd·s/m^2^, to capture the rod response. Subsequently, a 10 min light adaptation phase was initiated, followed by recording cone impulse responses at intensities of 3 cd·s/m^2^ and 10 cd·s/m^2^, as well as a 10 Hz flicker response. The ERG waveform data were assessed for amplitude and latency of response components, including flicker responses.

### Protein Analysis

For western blots, MEFs were lysed directly into denaturing gel loading buffer (Tris-HCl 125 mM pH6.8, glycerol 20% v/v, SDS 4% v/v, β-mercaptoethanol 10% v/v, bromophenol blue). Proteins were separated on Criterion TGX 7.5% 26 well precast gel (5671025, Biorad, Hercules CA USA) and electrophoretically transferred to Immobilon-FL 0.45 µm membrane (IPFL00005, Millipore, Burlington MA USA). Western blots were blocked with Licor Intercept (TBS) blocking buffer (927-60001, LI-COR, Lincoln NE USA) for 1 hr followed by primary antibody which was diluted into Intercept antibody diluent (927-66001, LI-COR) for 1 hr. Following primary antibody incubation, blots were washed 4 times in TBS and incubated for 1 hr with secondary antibody diluted in Intercept antibody diluent (927-66001, LI-COR) containing 0.01% SDS. After 4 washes in TBS, the blots were imaged with a LI-COR imager (Odyssey).

### Plasmids

Plasmids were assembled by TEDA assembly (Xia et al., 2019) into the pHAGE lentiviral backbone (Wilson et al., 2008). Mutations were generated by PCR amplification with mutated primers and the products TEDA assembled. All inserts were fully sequenced and matched NCBI reference sequence or expected mutant forms. Plasmids are listed in Table 1 and SnapGene files will be provided upon request.

**Table 1.** Plasmids.

| Name | Description* | Promot | Insert | Tags | Selection |
| --- | --- | --- | --- | --- | --- |
| GP801.1 | Flag-Ift43 | CMV | *M. musculus* Ift43 | 3xFlag | Bsd |
| GP1176.4 | Flag-Ift43<sup>E34K</sup> | CMV | *M. musculus* Ift43<sup>E34K</sup> | 3xFlag | Bsd |
| GP1177.10 | Flag-Ift43<sup>W172R</sup> | CMV | *M. musculus* Ift43<sup>W172R</sup> | 3xFlag | Bsd |
| GP1178.1 | Flag-Ift43<sup>ΔN127</sup> | CMV | *M. musculus* Ift43<sup>128-207</sup> | 3xFlag | Bsd |
| GP1181.1 | Flag-Ift43<sup>ΔC22</sup> | CMV | *M. musculus* Ift43<sup>1-184</sup> | 3xFlag | Bsd |
| GP1182.6 | Flag-Ift43<sup>ΔN21</sup> | CMV | *M. musculus* Ift43<sup>22-206</sup> | 3xFlag | Bsd |
| GP1183.1 | Flag-Ift43<sup>Δ127ΔC22</sup> | CMV | *M. musculus* Ift43<sup>128-184</sup> | 3xFlag | Bsd |

\*See S3 Data for the sequences of the proteins produced by these variants.

### Lentivirus Production

Lentiviral packaged pHAGE-derived plasmids (Wilson et al., 2008) were used for transfection. These vectors are packaged by a third-generation system comprising four distinct packaging vectors (Tat, Rev, Gag/Pol, VSV-g) using HEK 293T cells as the host. DNA (Backbone: 5µg; Tat: 0.5 µg; Rev: 0.5 µg; Gag/Pol: 0.5 µg; VSV-g: 1 µg) was delivered to the HEK cells using calcium phosphate precipitates. After 48 hrs, the supernatant was harvested, filtered through a 0.45 mm PES filter (Sigma-Aldrich, SLHPR33RS), and added to ∼50% confluent cells. After 24 hrs, cells were selected with blasticidin (60 µg/ml).

### Statistics

Data groups were analyzed by as described in the figure legends using GraphPad Prism (San Diego CA USA). Differences between groups were considered statistically significant if p < 0.05. Statistical significance is denoted with ****p≤0.0001, ***p≤0.001, **p≤0.01, *p≤0.05, or ns not significant. Error bars are standard deviations (SD) unless otherwise marked. Solid bars in violin plots indicate the median, doted lines indicate quartiles.

## Supporting information

S1 Data. Raw data underlying Fig 1

S2 Data. Diagram of Alleles and Genotyping Details

S3 Data. Sequence of Ift43 deletions

S4 Data. Necropsy, CT data, and EFIC phenotyping

S5 Data. Raw data underlying the graphs in Fig 5-10

S6 Data. Uncropped western blots

S7 Data. ImageJ Macros

S1 Figure. OPT images of Ift43 mutants

S2 Figure. OPT image to illustrate malformed somites.

S3 Figure. Ift43 mutant lymphatic cilia do not label with Arl13b antibodies

S4 Figure. Ift43 KO causes defects in mouse embryonic lymphatic vessel development

S5 Figure. Retinal protein distribution is not greatly affected by the loss of Ift43

S6 Figure. Arl13b requires Ift43 for ciliary entry

## Acknowledgments

Lymphatic analysis used resources provided by the South Dakota State University Functional Genomics Core Facility (RRID:SCR_023786) supported in part by the National Science Foundation/EPSCoR Grant No. 0091948, the South Dakota Agricultural Experiment Station, and by the State of South Dakota.

Flow Cytometry Resources at UMass Chan were supported by National Institutes of Health NIH S10OD028576.

Mouse strain Ift43^tm1a(EUCOMM)Hmgu^ was generated at the Baylor College of Medicine as part of the Baylor College of Medicine, Sanger Institute, and MRC Harwell (BaSH) Consortium for the NIH Common Fund program for Knockout Mouse Production and Cryopreservation (1U42RR033192-01) and Knockout Mouse Phenotyping (1U54HG006348-01).

## Ethics Statement

Mouse research was carried out at the University of Massachusetts Chan Medical School with IACUC approval (PROTO201900265). This IACUC follow the regulations of US Department of Agriculture Animal Welfare Act and the standards/principles of the Public Health Service Policy on Humane Care and Use of Laboratory Animals, AVMA Guidelines on Euthanasia, U.S. Government Principles for the Utilization and Care of Vertebrate Animals Used in Testing, Research and Training, and the Guide for the Care and Use of Laboratory Animals.

## Abbreviations

AVSD: atrioventricular septal defect
BSA: bovine serum albumin
CT: computed tomography
DAPI: 4′,6-diamidino-2-phenylindole
DEXV: dextroversion
EFIC: episcopic fluorescence image capture
E: embryonic day
ERG: electroretinograms
G: Golgi complex
IS: inner segment
IFT: intraflagellar transport
INL: inner nuclear layer
LA: left atria
LV: left ventricle
mVSD: muscular ventricular septal defect
OS: outer segment
ONL: outer nuclear layer
OPL: outer plexiform layer
P: postnatal day
PTA: persistent truncus arteriosus
RA: right atria
ROI: region of interest
RV: right ventricle
SIf: superior-inferior ventricles
TEM: transmission electron microscopy
WGA: Wheat germ agglutinin

## Supplemental Files

**S1 Figure. OPT images of Ift43 mutants**

i-iii) OPT images of *Ift43* mutants with the planes marked.

i’-iii’) Single planes from i-iii to illustrate medial mandibular seam (i’), a hinge blister (ii’), and medial oral cleft (iii’). Scale bar =.

**S2 Figure. OPT image to illustrate malformed somites.**

Surface rendered OPT image of an *Ift43* mutant to illustrate the malformed somites (arrows). Scale bar = _.

**S3 Figure. Ift43 mutant lymphatic cilia do not label with Arl13b antibodies**

Images of control and experimental E15.5 skin stained with Nrp2 (red) Prox1 (grey) and Ar13b (grey). Scale bar = 20 µm, images are maximum projections of stacks taken at 0.5 µm intervals. Note the lack of Arl13b-positive cilia in the experimental tissue.

**S4 Figure. Ift43 KO causes defects in mouse embryonic lymphatic vessel development**

Example of data used for quantification of proliferation. Lymphatic vessels labeled via Nrp2 and Prox1. Proliferating cells labeled via phospho-histone H3 (pHH3). White arrows highlight colocalization of Prox1 and pHH3, indicating proliferating lymphatic endothelial cells. Scale bar = 100 µm, images are maximum projections of stacks taken at 1.0 µm intervals.

**S5 Figure. Retinal protein distribution is not greatly affected by the loss of Ift43**

Confocal images of agarose-embedded retinal cross-sections. P25 control (*Ift43^flox/+^/iCre75)* and experimental (*Ift43^flox/flox^/iCre75)* littermate mice stained with antibodies against outer segment proteins Prom1 (A, green) and Rom1 (B, green) an inner segment protein syntaxin-3 (C, red). Panel D shows colocalization of syntaxin 3 (red) with acetylated tubulin (green). Nuclei are labelled with DAPI (4′,6-diamidino-2-phenylindole) in blue. Images are maximum intensity z-projections of 20 images taken at 0.7 µm intervals. OS: outer segment; IS: inner segment; ONL: outer nuclear layer, OPL: outer plexiform layer; INL: inner nuclear layer. Scale bars = 20 µm (A-C) or 10 µm (D). Protein intensity profiles of Prom1, Rom1 (green), and syntaxin-3 (red) along white lines shown at right. n=3 animals examined.

**S6 Figure. Arl13b requires Ift43 for ciliary entry**

**(A)** Cilia stained for acetylated α-tubulin (AcTub, red) and Arl13b (green [left], grey [right]) in wild type (10640.4), *Ift43* mutant (27260.1), and *Ift43* mutant fibroblasts rescued with wild type Ift43 tagged with Flag at the N-terminus (GP801). Images are maximum projections of stacks taken at 0.14 µm intervals. Scale bars = 1 µm.

**(B)** Quantification of ciliary Arl13b in wild type (10640.4), *Ift43* mutant (27260.1), and *Ift43* mutant fibroblasts transfected with a wild-type or mutant constructs described in Fig 7A. The quantification tools of CiliaQ were used to measure fluorescence intensity. n>100 for each condition. Violin plots show median (bold dashed line) and quartiles (dashed lines). Note that some outliers are not plotted but all data was used for statistical analysis and is available in S5 Data. ****p≤0.0001, **p≤0.01, *p≤0.05, ns not significant as compared to wild-type (#, top row), *ift43* mutant (# middle row) or wild-type rescue (#, bottom row) by one-way ANOVA with Tukey’s multiple comparisons post-hoc test.

**(C)** Quantification of ciliary Arl13b in *Ift140* MEFs. Parameters are the same as described in **(B)**. Raw data underling the graph is available in S5 Data.

**S1 Table.**
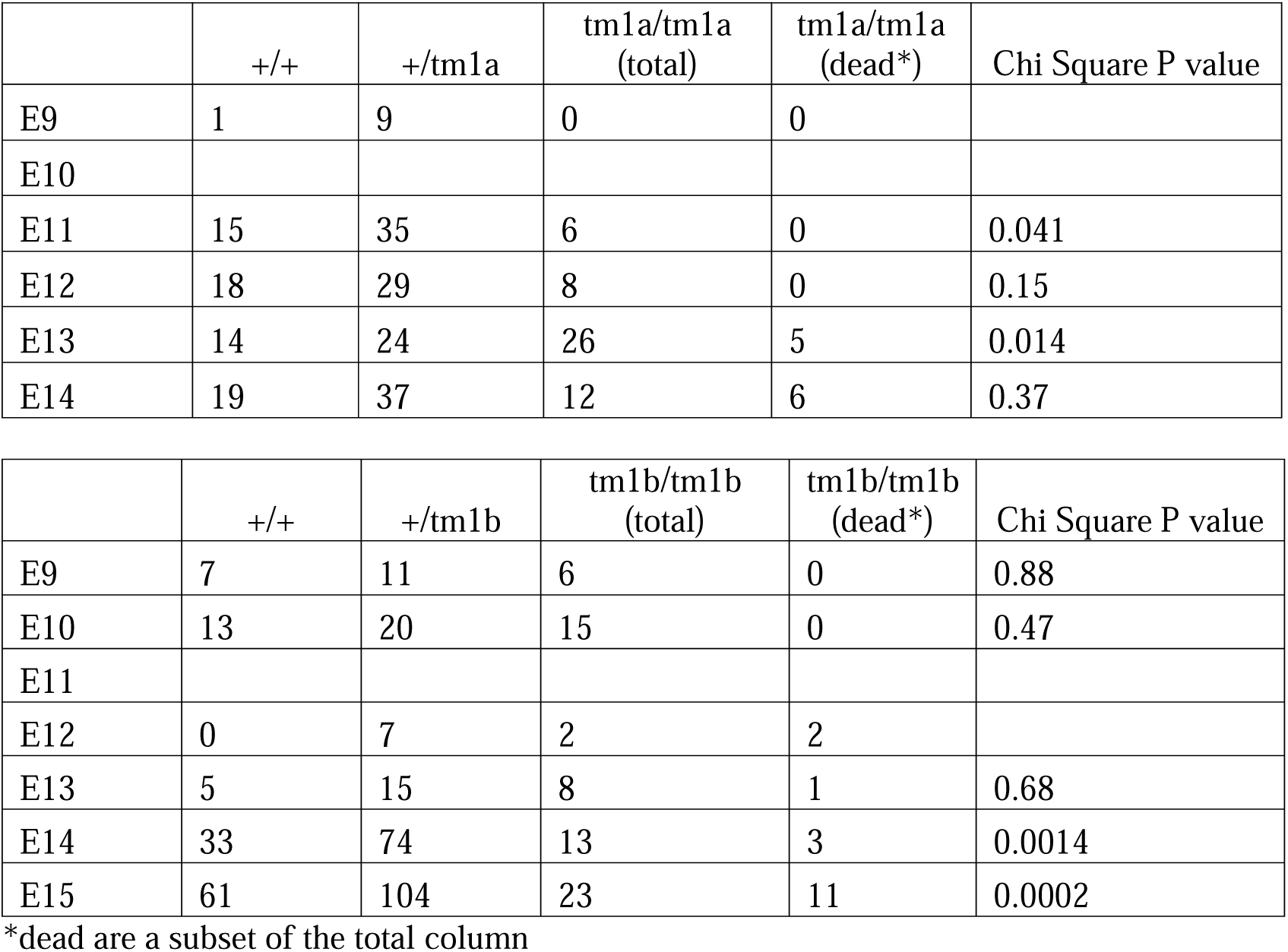
Embryo Genotype Distribution.

**S2 Table.** Antibodies.

| **Antibody (antigen species)** | **Species Raised In** | **Clone or Antibody Name (Isotype)** | **Supplier** | **Dilution** |
| --- | --- | --- | --- | --- |
| **Primaries** |  |  |  |  |
| Flag | Mouse | M2 (IgG1) | F1804, Sigma-Aldrich, St. Louis MO USA | 1:1000 |
| Arl13b (mouse) | Mouse | N295B/66 (IgG2a) | NeuroMab, Davis CA USA | 1:1000 |
| Arl13b (mouse) | Rabbit |  | 17711-1-AP, Proteintech, Rosemont IL USA | 1:1000 |
| Gli1 (human, mouse) | Rabbit | V812 | 2534, Cell Signaling, Danvers MA USA | 1:500 |
| Gli2 (mouse) | Goat | AF3635 | R&D Systems, Minneapolis MN USA | 1:1000 |
| Gli3 (human) | Goat | AF3690 | R&D Systems | 1:200 |
| Smo (human) | Mouse | E5 (IgG2a) | sc-166685, Santa Cruz Biotechnology, Dallas TX USA | 1:100 |
| Ift27 (mouse) | Rabbit | 719 | (Keady et al., 2012) | 1:250 |
| Ift88 (mouse) | Rabbit | 448 | (Pazour et al., 2002) | 1:250 |
| BBS5 (human) | Mouse | B-11 (IgG1) | sc-515331, Santa Cruz Biotechnology | 1:500 |
| ~~BBS2 (human)~~ | ~~Mouse~~ | ~~A-12 (IgG1)~~ | ~~sc-365355, Santa Cruz Biotechnology~~ | ~~1:100~~ |
| ~~BBS2 (human)~~ | ~~Rabbit~~ |  | ~~11188-2 AP, Proteintech~~ | ~~1:500~~ |
| ~~LZTFL1 (human)~~ | ~~Rabbit~~ |  | ~~17073-1 AP, Proteintech~~ | ~~1:500~~ |
| α-Tubulin (sea urchin) | Mouse | B-5-1-2 (IgG1) | T6074, Sigma-Aldrich | 1:5000 |
| GAPDH (human) | Rabbit | 14C10 | 3683S, Cell Signaling, Danvers MA USA | 1:5000 |
| Acetylated α-Tubulin (sea urchin) | Mouse | 6-11B-1 (IgG2b) | T7451, Sigma-Aldrich | 1:10000 |
| **Secondaries** |  |  |  |  |
| Rabbit IgG-Alexa 488 | Goat |  | A11034, Invitrogen, Waltham MA USA | 1:1000 |
| Rabbit IgG-Alexa 568 | Goat |  | A11011, Invitrogen | 1:1000 |
| Rabbit IgG-Alexa 594 | Donkey |  | A21207, Invitrogen | 1:1000 |
| Goat IgG-Alexa 594 | Donkey |  | A11058, Invitrogen | 1:1000 |
| Mouse IgG-Alexa 488 | Goat |  | A11017, Invitrogen | 1:1000 |
| Mouse IgG-Alexa 488 | Donkey |  | A32766, Invitrogen | 1:1000 |
| Mouse IgG-Alexa 568 | Donkey |  | A10037, Invitrogen | 1:1000 |
| Mouse IgG-Alexa 594 | Goat |  | A11032, Invitrogen | 1:1000 |
| Mouse IgG1-Alexa 488 | Goat |  | A21121, Invitrogen | 1:1000 |
| Mouse IgG1-Alexa 568 | Goat |  | A21124, Invitrogen | 1:1000 |
| Mouse IgG2a-Alexa 488 | Goat |  | A21131, Invitrogen | 1:1000 |
| Mouse IgG2a-Alexa 568 | Goat |  | A21134, Invitrogen | 1:1000 |
| Mouse IgG2b-Alexa 488 | Goat |  | A21141, Invitrogen | 1:1000 |
| Mouse IgG2b-Alexa 568 | Goat |  | A21144, Invitrogen | 1:1000 |
| Rabbit IRDye 800CW | Goat |  | 925-32211 LI-COR | 1:20,000 |
| Mouse IRDye 680RD | Donkey |  | 925-68072 LI-COR | 1:20,000 |
| Goat IRDye 800CW | Donkey |  | 925-32214 LI-COR | 1:20,000 |

**S3 Table.**
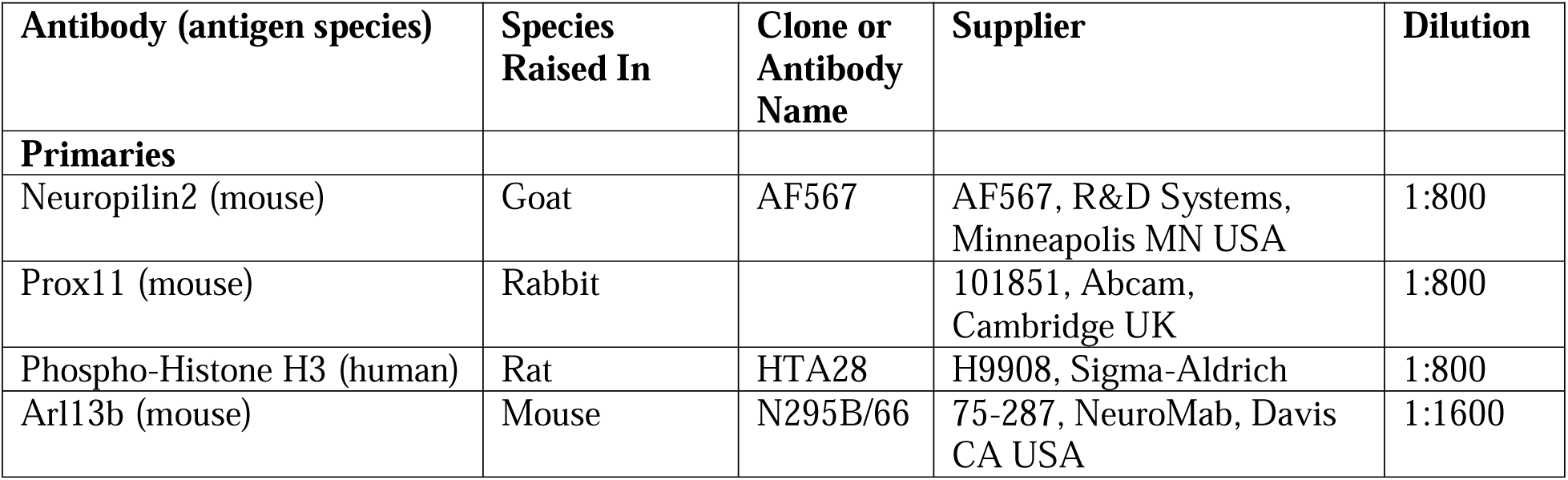
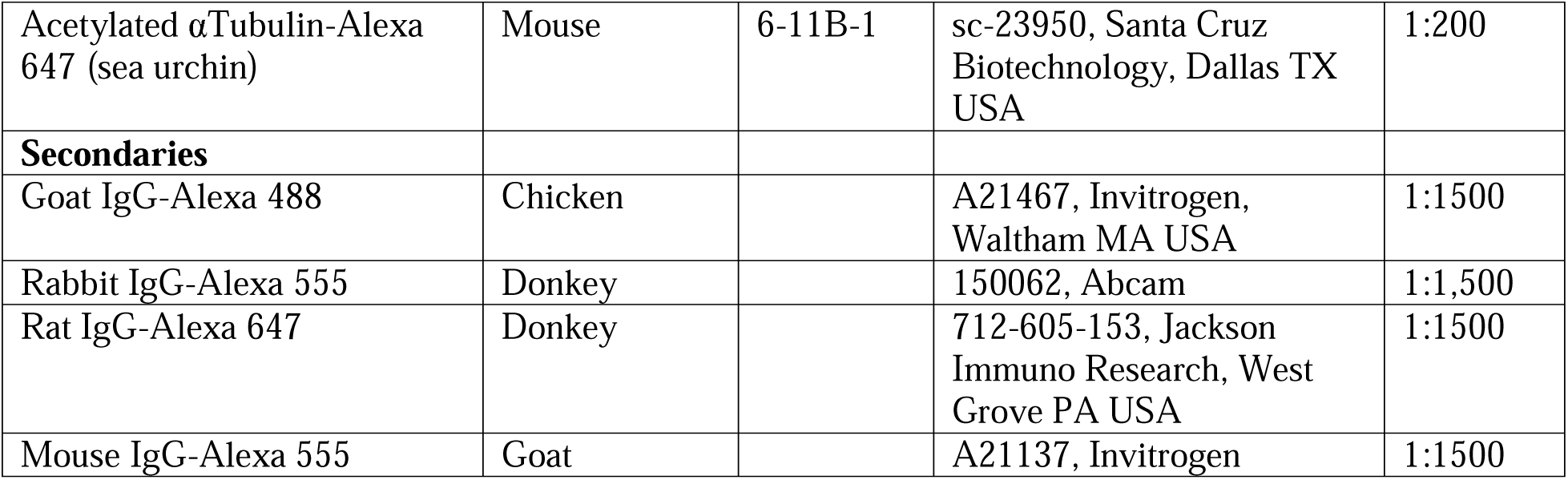
Antibodies used for lymphatic analysis.

| Antibody (antigen species) | Species Raised In | Clone or Antibody Name | Supplier | Dilution |
| --- | --- | --- | --- | --- |
| **Primaries** |  |  |  |  |
| Neuropilin2 (mouse) | Goat | AF567 | AF567, R&D Systems, Minneapolis MN USA | 1:800 |
| Prox11 (mouse) | Rabbit |  | 101851, Abcam, Cambridge UK | 1:800 |
| Phospho-Histone H3 (human) | Rat | HTA28 | H9908, Sigma-Aldrich | 1:800 |
| Arl13b (mouse) | Mouse | N295B/66 | 75-287, NeuroMab, Davis CA USA | 1:1600 |
| Acetylated αTubulin-Alexa 647 (sea urchin) | Mouse | 6-11B-1 | sc-23950, Santa Cruz Biotechnology, Dallas TX USA | 1:200 |
| **Secondaries** |  |  |  |  |
| Goat IgG-Alexa 488 | Chicken |  | A21467, Invitrogen, Waltham MA USA | 1:1500 |
| Rabbit IgG-Alexa 555 | Donkey |  | 150062, Abcam | 1:1,500 |
| Rat IgG-Alexa 647 | Donkey |  | 712-605-153, Jackson Immuno Research, West Grove PA USA | 1:1500 |
| Mouse IgG-Alexa 555 | Goat |  | A21137, Invitrogen | 1:1500 |

**S1 Data. Raw data underlying Fig 1**

Blast values and alignments used to generate Fig 1.

**S2 Data. Diagram of Alleles and Genotyping Details**

**(A)** Diagram of the alleles used in this study. The start codon is in exon 2 with the termination codon in exon 9. Frt and LoxP recombination sites are represented by circles and arrows respectively. Exons and introns are not drawn to scale and the introns do not reflect size changes that occur during recombination.

**(B)** Primers and PCR product sizes from the various alleles.

**(C)** Sequence of wild type Ift43 and the predicted protein after Cre excision of the flox allele.

**S3 Data. Sequence of Ift43 deletions**

Sequence of the various Ift43 variants used in this work.

**S4 Data. Necropsy, CT data, and EFIC phenotyping**

Phenotyping data behind Figs 3A-3M and 4.

**S5 Data. Raw data underlying the graphs in Fig 5-10**

Raw data and statistical analysis used to generate the graphs in Figs 5-10.

**S6 Data. Uncropped western blots**

Uncropped western blots used for figures and quantification.

**S7 Data. ImageJ Macros**

Code to use segmentations created by ilastik (Berg et al., 2019) to create image stacks that can be analyzed by CiliaQ (Hansen et al., 2021).

**S1 Video. OPT head render**

OPT surface rendering of a control and *Ift43* mutant head illustrating the malformed face and head seen in the mutants.

**S2 Video. OPT whole body render**

OPT surface rendering of an *Ift43* mutant embryo showing the malformed face and head, the body wall defect, and distorted tail.

