## Supplementary material for "Ift43 Controls the Ciliary Levels of Gli2 and Gli3": S2 Data. Diagram of Alleles and Genotyping Details

**A**

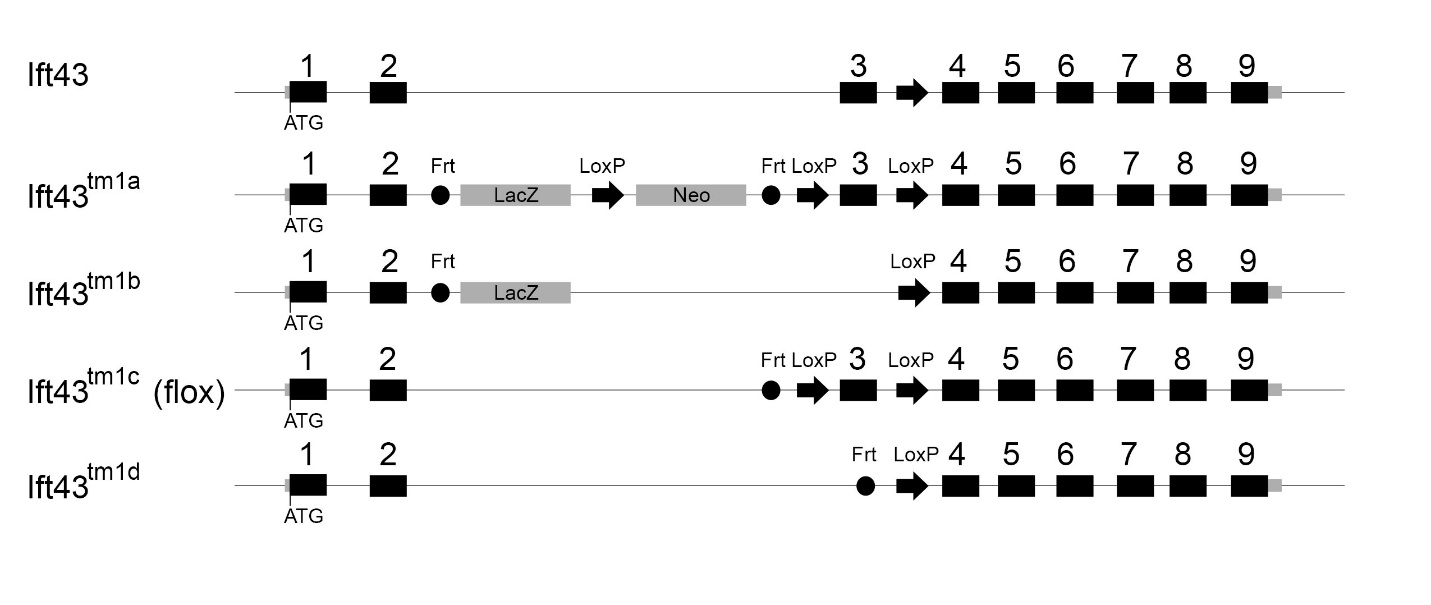

**B**

Genotyping:

37736-LacF + 37736-R	Wild Type	no product

Ift43^tm1a^	4223 bp

Ift43^tm1b^	657 bp

Ift43^tm1d^	no product

37736-F + 37736-TTR	Wild Type	307 bp

Ift43^tm1a^	7386 bp

Ift43^tm1c^ (Flox)	482 bp

37736-NeoF + 37736-TTR	Wild Type	no product

Ift43^tm1a^	563 bp

37736-LacF	GCTACCATTACCAGTTGGTCTGGTGTC	Tm = 64

37736-neoF	GGGATCTCATGCTGGAGTTCTTCG	Tm = 62

37736-loxF	GAGATGGCGCAACGCAATTAATG	Tm = 58

37736-TTR	GCTCTTAACTGCTGACCTATCTCTCC	Tm = 62

37736-R	GTGTGATTCCAGAACACTGCTGATC	Tm = 61

37736-F	CATCGGTGCTTCGAATTCATGG	Tm = 58
