## Supplementary material for "Ift43 Controls the Ciliary Levels of Gli2 and Gli3": S3 Data. Sequence of Ift43 deletions

>GP801.1 Ift43^WT^ 1-206

3xFlag-MDDLLDLGEERRRSSATSGAKMGRRAHQESTQAENYLNSKNSSLTQTGEAPPPKPPRRQ GGWADDSMKMAKLGKKVSEEMEDHRLRQQNLMGSDDDDIPVIPDLEDVQEEDFVLQVASPPSIQVNRVMTYRDLDNDLMKYSAFQTLDGEIDLKLLTKVLAPEHEVREDDVGWDWDHLYTEVSSELLTEWDLLRAEKEDPVGPSRYI*

>GP1176.4 Ift43^E34K^

3xFlag-MDDLLDLGEERRRSSATSGAKMGRRAHQESTQA**K**NYLNSKNSSLTQTGEAPPPKPPRRQ GGWADDSMKMAKLGKKVSEEMEDHRLRQQNLMGSDDDDIPVIPDLEDVQEEDFVLQVASPPSIQVNRVMTYRDLDNDLMKYSAFQTLDGEIDLKLLTKVLAPEHEVREDDVGWDWDHLYTEVSSELLTEWDLLRAEKEDPVGPSRYI*

>GP1177.10 Ift43^W172R^

3xFlag-MDDLLDLGEERRRSSATSGAKMGRRAHQESTQAENYLNSKNSSLTQTGEAPPPKPPRRQ GGWADDSMKMAKLGKKVSEEMEDHRLRQQNLMGSDDDDIPVIPDLEDVQEEDFVLQVASPPSIQVNRVMTYRDLDNDLMKYSAFQTLDGEIDLKLLTKVLAPEHEVREDDVG**R**DWDHLYTEVSSELLTEWDLLRAEKEDPVGPSRYI*

>GP1178.1 Ift43^ΔN127^ (128-206)

3xFlag-MTYRDLDNDLMKYSAFQTLDGEIDLKLLTKVLAPEHEVREDDVGWDWDHLYTEVSSE LLTEWDLLRAEKEDPVGPSRYI*

>GP1181.1 Ift43^ΔC22^ (1-184)

3xFlag-MDDLLDLGEERRRSSATSGAKMGRRAHQESTQAENYLNSKNSSLTQTGEAPPPKPPRRQ GGWADDSMKMAKLGKKVSEEMEDHRLRQQNLMGSDDDDIPVIPDLEDVQEEDFVLQVASPPSIQVNRVMTYRDLDNDLMKYSAFQTLDGEIDLKLLTKVLAPEHEVREDDVGWDWDHLYTEVSSE*

>GP1182.6 Ift43^ΔN21^ (22-206)

3xFlag-MGRRAHQESTQAENYLNSKNSSLTQTGEAPPPKPPRRQGGWADDSMKMAKLGKKVSE EMEDHRLRQQNLMGSDDDDIPVIPDLEDVQEEDFVLQVASPPSIQVNRVMTYRDLDNDLMKYSAFQTLDGEIDLKLLTKVLAPEHEVREDDVGWDWDHLYTEVSSELLTEWDLLRAEKEDPVGPSRYI*

>GP1183.1 Ift43^ΔN127ΔC22^ (128-184)

3xFlag-MTYRDLDNDLMKYSAFQTLDGEIDLKLLTKVLAPEHEVREDDVGWDWDHLYTEVSSE*

>post flox deletion 1-44 with 6 nonsense amino acids before termination

MDDLLDLGEERRRSSATSGAKMGRRAHQESTQAENYLNSKNSSLTQTGE**IGKESF***
