## Supplementary material for "Ift43 Controls the Ciliary Levels of Gli2 and Gli3": S6 Data. Uncropped western blots

Supplemental Data 6. Uncropped Western Blots
Related to Figure 9 Aa-Ab

Blot Used for Quantification Only

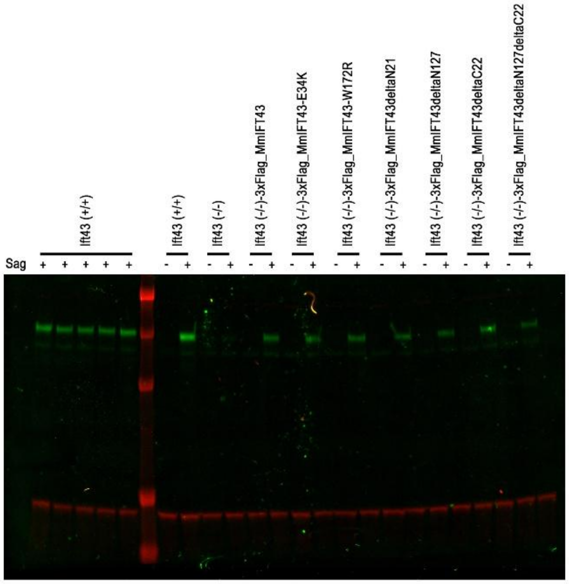


Cropped blot blot Used in Figure 9

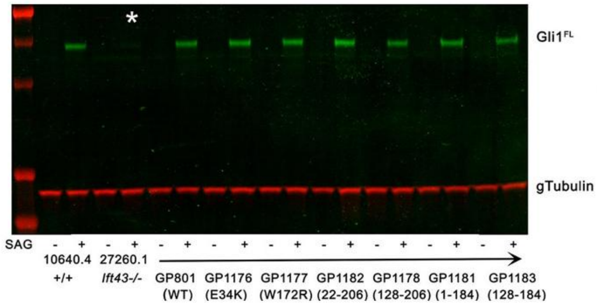


Blot Used for Quantification Only

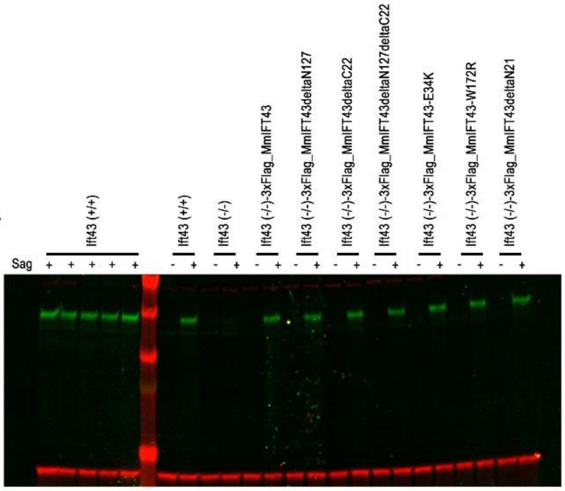


Blot Used for Figure and Quantification
Region cropped indicated by white box

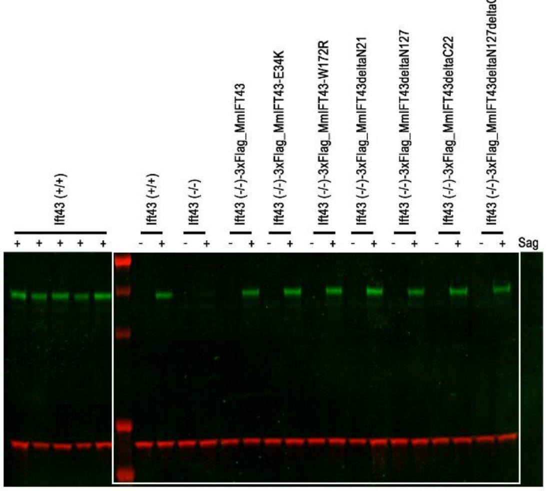

Supplemntal Data 6. Uncropped Western Blots Related to Figure 9 Ca-Cc

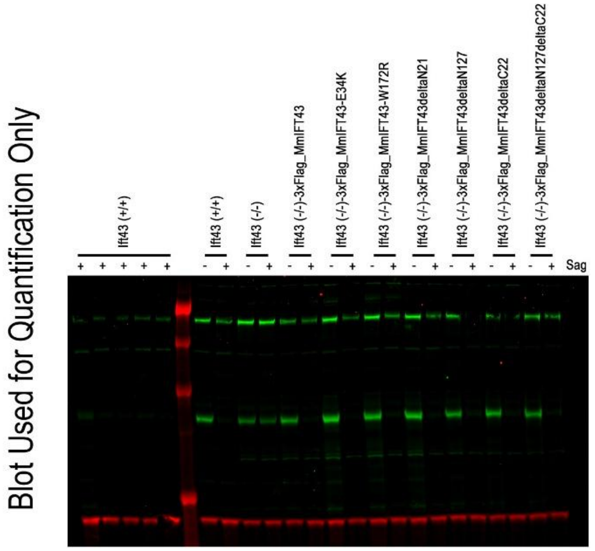


Ca Cropped blot blot Used in Figure 9

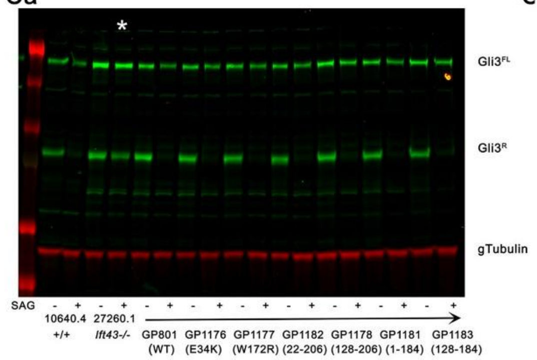


C

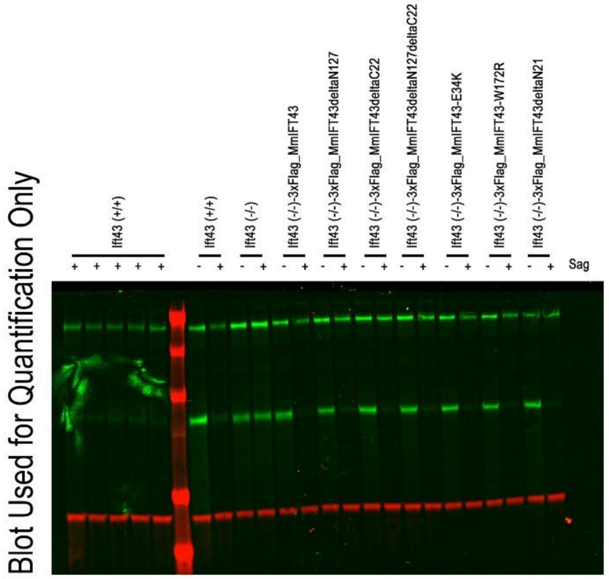


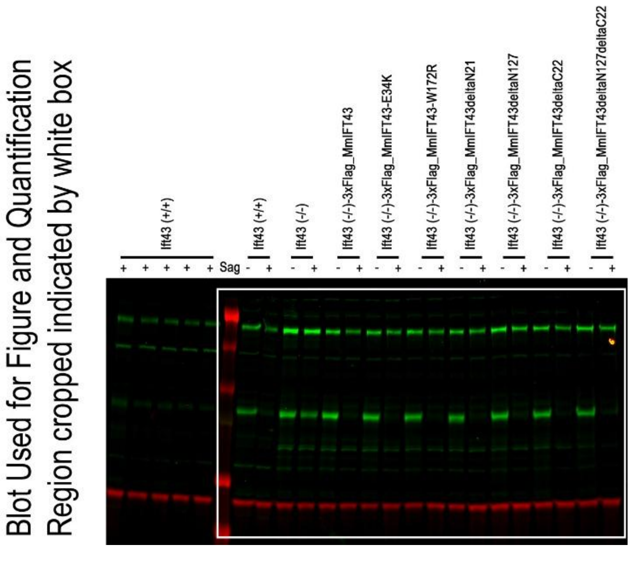

Supplemental Data. Uncropped Western Blots
Related to Figure 9 Ba-Bc

Blot Used for Figure and Quantification
Region cropped indicated by white box

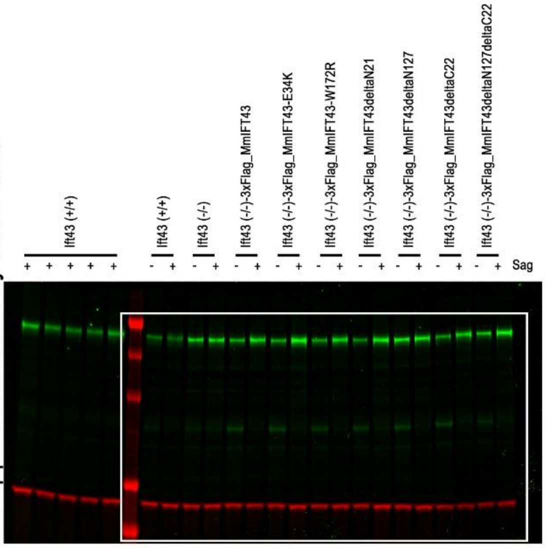


**Ba** Cropped blot blot Used in Figure 9

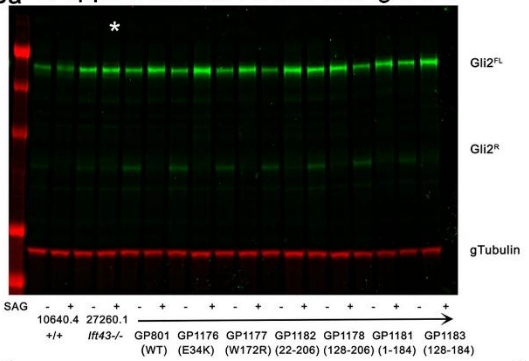


Blot Used for Quantification Only

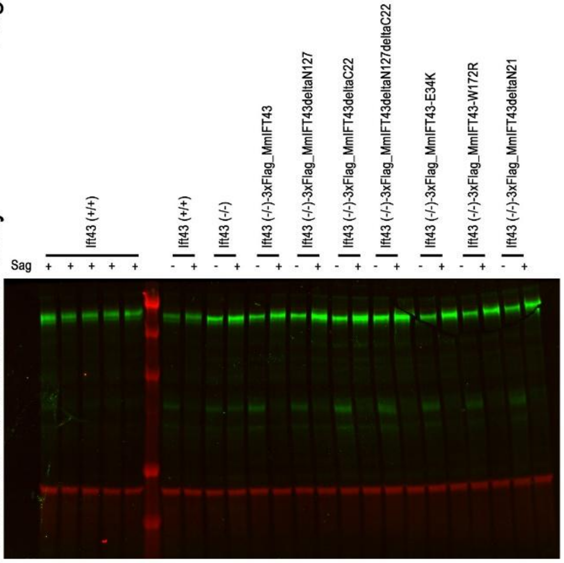


Blot Used for Quantification Only

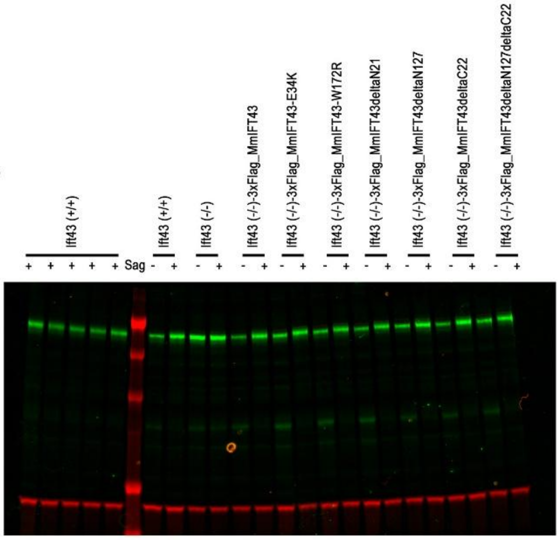
