## Supplementary material for "Ift43 Controls the Ciliary Levels of Gli2 and Gli3": S7 Data. ImageJ Macros

**The following code converts the pixel classification performed using the simple segmentation funtion of ilastik, in which pixels are classified as cilia (value=1) and background (value=2), to a binary mask using imageJ.**

**To run, open the ilastik simple segmentation images in ImageJ, load the code into the script window, and run repeatedly until all the open images are closed.**

**Note: the scaling is hardcoded sso it may need to be modified to match your imaging parameters.**

//get number of slices in image

stackheight = (nSlices);

// write the number of slices to the log

write(stackheight);'

// reset the properties of the file that were stripped out by iLastik and convert to 16 bit image

run("Convert to Mask", "background=Light black");

run("Properties...", "channels=1 slices=stackheight frames=1 pixel_width=0.1559815 pixel_height=0.1559815 voxel_depth=0.37");

setOption("ScaleConversions", true);

run("16-bit");

// Get the current image title

originalTitle = getTitle();

// Specify the part of the name to replace and the replacement text

oldText = "_Simple Segmentation_.tiff";

newText = "_mask";

// Replace the old part of the file name with the new text

newTitle = replace(originalTitle, oldText, newText);

// Get the directory of the original file

directory = getDirectory("current");

// Create the full path for the new file

newFilePath = directory + newTitle;

// Save the current image with the new name

saveAs("tif", newFilePath); // Change "jpeg" to your desired format (e.g., "png", "tiff")

// Close the image without any dialogue

close();

**The following code inserts the ilastik binary masks created with the previous code into the image stacks producing an image stack in the format needed by CiliaQ.**

**The code will prompt for a file directory containing the image stacks and the coresponding masks generated with previous code. These will be combined to produce a three channel file that is ready to be fed into ciliaQ.**

//getImageID();

close("*"); // closes any open windows. can be commented out if you don't want the windows to close.

//selectImage(-2181);

setBatchMode(true);

// Prompt the user to select a directory

dir = getDirectory("Select a Directory");

// Get a list of all TIFF files in the directory

fileList = getFileList(dir);

// Loop through the list of files

for (i = 0; i < fileList.length; i++) {

fileName = fileList[i];

// Check for File A (_mask.tif) and extract root name

if (indexOf(fileName, "_mask.tif") >=0) {

// Extract the root name (without "_mask.tif")

rootName = substring(fileName, 0, lengthOf(fileName) - 9);

// Construct the corresponding File B name

fileBName = rootName + ".tif"; // Assuming File B ends with ".tif"

// Check if File B exists in the directory

found = false;

for (j = 0; j < fileList.length; j++) {

if (fileList[j] == fileBName) {

found = true;

break;

}

}

if (found) {

// Open File A (channel 1)

open(dir + fileName);

channel1ID = (getImageID() -1); // Get the ID of channel 1

numSlicesA = (nSlices); // Get number of slices in channel A

run("Stack to Images"); // Split File A into separate images // added

write (fileName);

write (channel1ID);

write (fileBName);

write (numSlicesA);

// Open File B (channels 2 and 3)

open(dir + fileBName);

channel2ID = (getImageID() - 1); // Get the ID of channel 2

run("Stack to Images"); // Split File B into separate images

//channel3ID = (getImageID() + numSlicesA); // Get the ID of channel 3

//close(); // Close the original two-channel image

// Create a new hyperstack to hold the combined channels

newImage("Combined Hyperstack", "16-bit", getWidth(), getHeight(), 3, numSlicesA, 1);

hyperstackID = getImageID();

// Loop through each slice and copy the corresponding layers to the new hyperstack

for (j = 0; j < numSlicesA; j++) {

curr_slice = (j * 3) + 1;

// Copy slice from channel 1 (File A) to channel 1

selectImage(channel1ID - j);

//setSlice(j);

run("Copy");

selectImage(hyperstackID);

setSlice(curr_slice); // Set slice for channel 1 in the new hyperstack

run("Paste");

curr_image = j * 2;

// Copy slice from channel 2 (File B) to channel 2

selectImage(channel2ID - curr_image);

//setSlice(j);

run("Copy");

selectImage(hyperstackID);

setSlice(curr_slice + 1); // Set slice for channel 2 in the new hyperstack

run("Paste");

// Copy slice from channel 3 (File B) to channel 3

selectImage(channel2ID - curr_image - 1);

//setSlice(j);

run("Copy");

selectImage(hyperstackID);

setSlice(curr_slice + 2); // Set slice for channel 3 in the new hyperstack

run("Paste");

}

//run("Channels Tool...");

Stack.setChannel(1);

run("Grays");

Stack.setChannel(2);

run("Red");

Stack.setChannel(3);

run("Green");

// Construct the new filename

newFileName = dir + rootName + "_3channel.tif";

// To set the proper scale

run("Properties...", "channels=3 slices=numSlicesA frames=1 pixel_width=0.1559815 pixel_height=0.1559815 voxel_depth=0.37");

// Save the new hyperstack

saveAs("Tiff", newFileName);

// Close the combined hyperstack

close();

close(channel1ID);

close(channel2ID);

//close(channel3ID);

} else {

// Warning if corresponding File B not found

showMessage("Warning", "Corresponding file for " + fileName + " not found.");

}

}

}

write("DONE");

// Function to check if a string ends with a specified suffix

function endsWith(str, suffix) {

return substring(str, lengthOf(str) - lengthOf(suffix), lengthOf(str)) == suffix;

}

// Function to get the length of a string

function lengthOf(str) {

return str.length;

}
