## Supplementary figures and images for "Ift43 Controls the Ciliary Levels of Gli2 and Gli3"

### S1 Figure. OPT images of Ift43 mutants

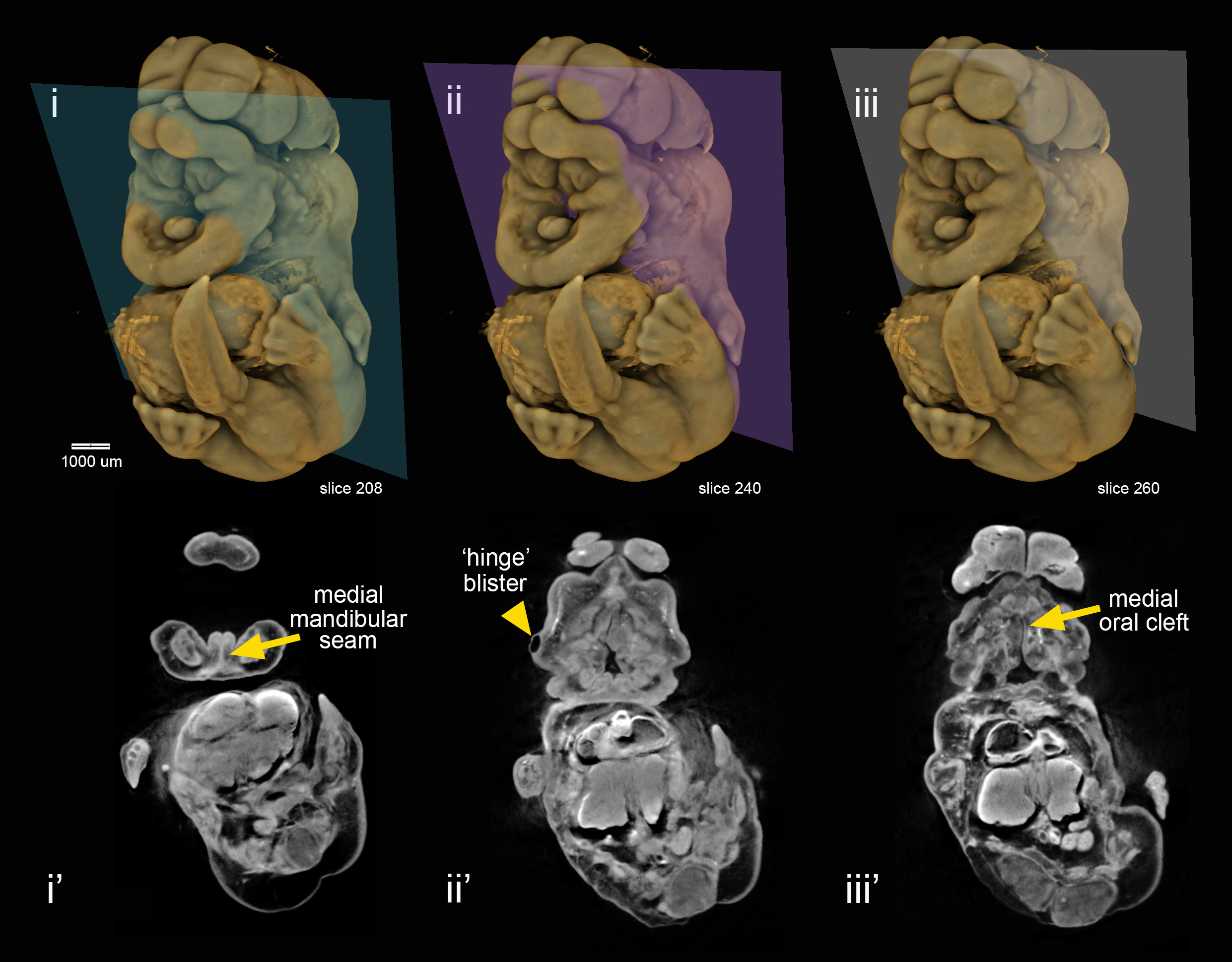

### S2 Figure. OPT image to illustrate malformed somites.

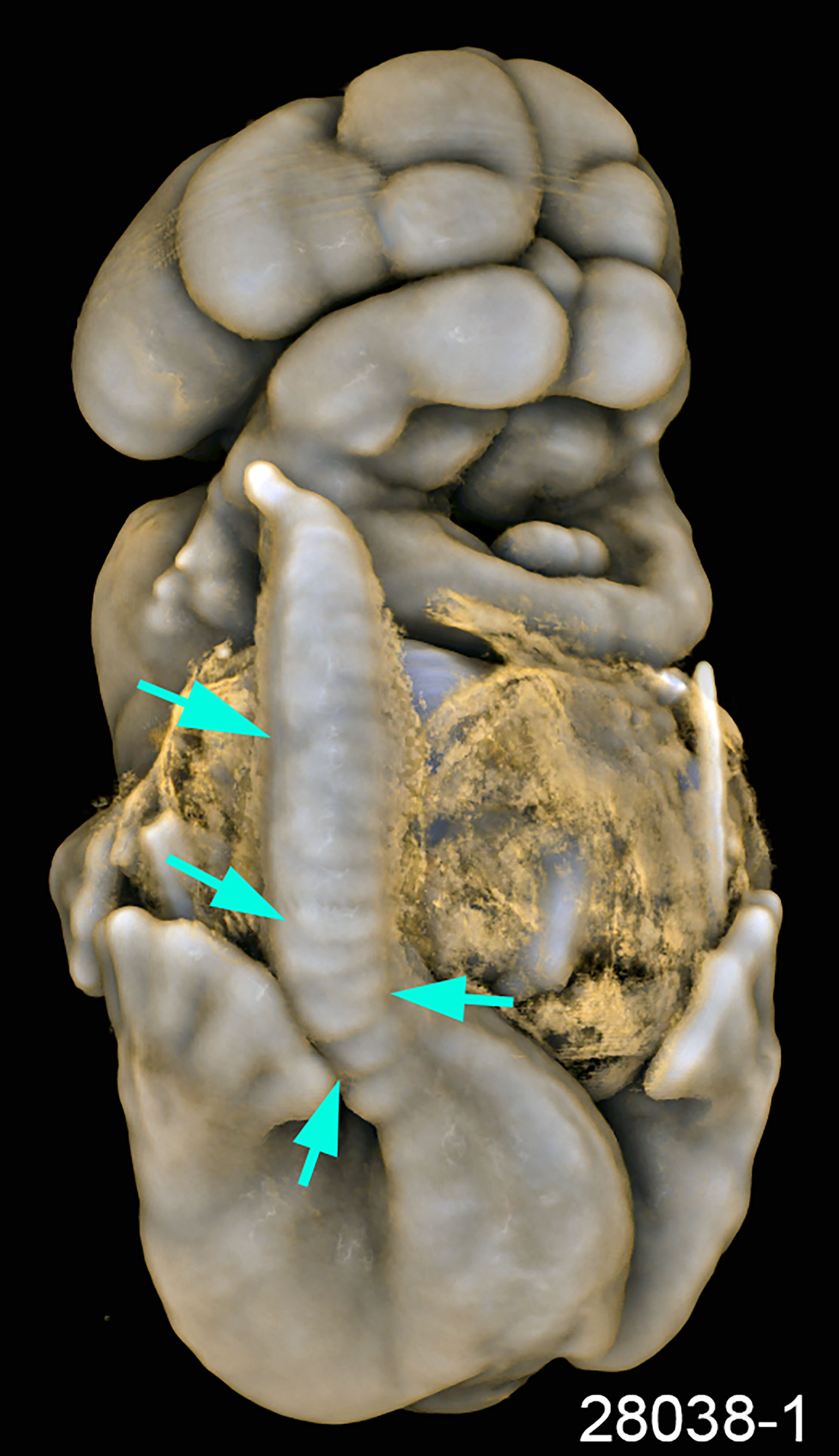

### S3 Figure. Ift43 mutant lymphatic cilia do not label with Arl13b antibodies

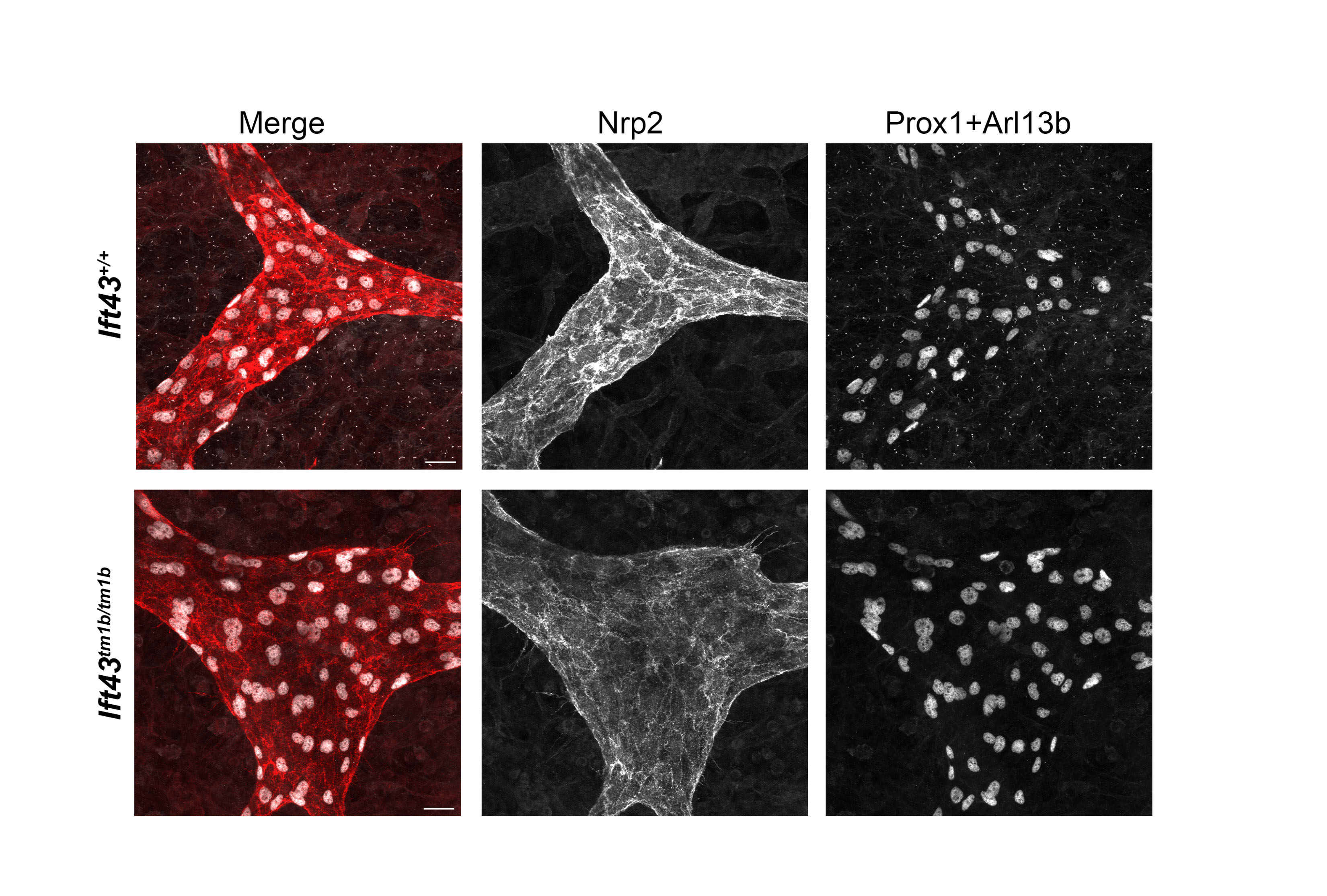

### S4 Figure. Ift43 KO causes defects in mouse embryonic lymphatic vessel development

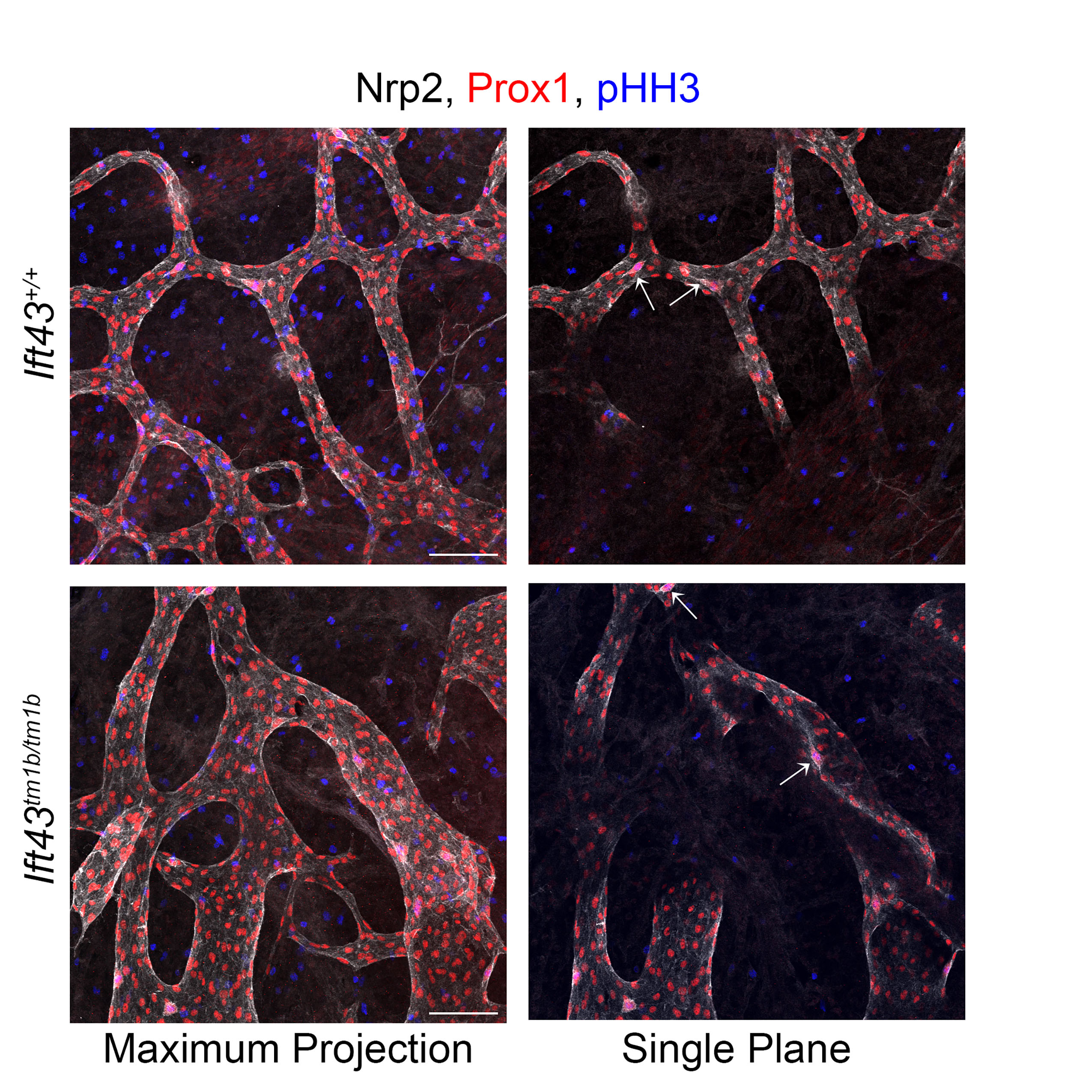

### S5 Figure. Retinal protein distribution is not greatly affected by the loss of Ift43

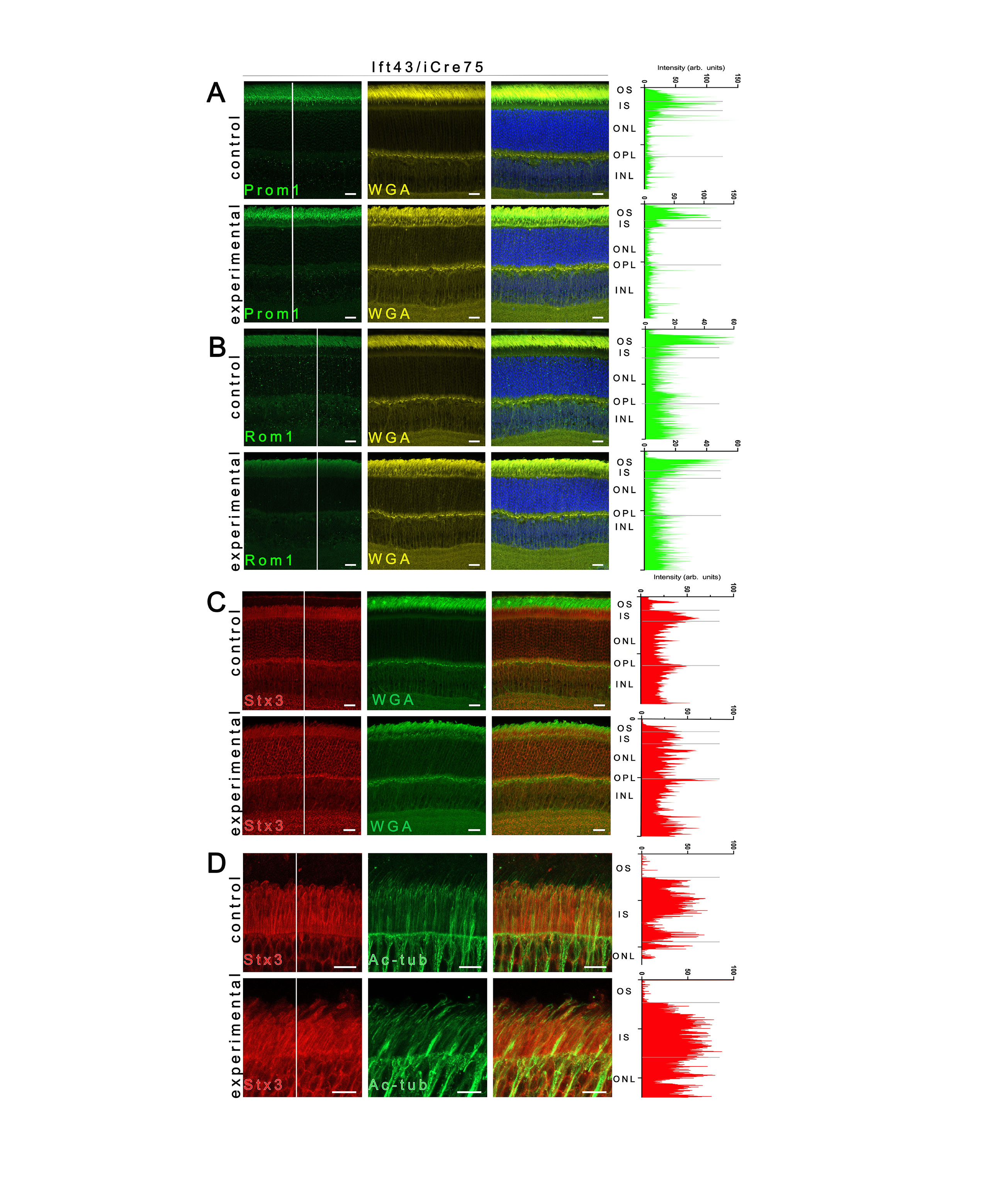

### S6 Figure. Arl13b requires Ift43 for ciliary entry

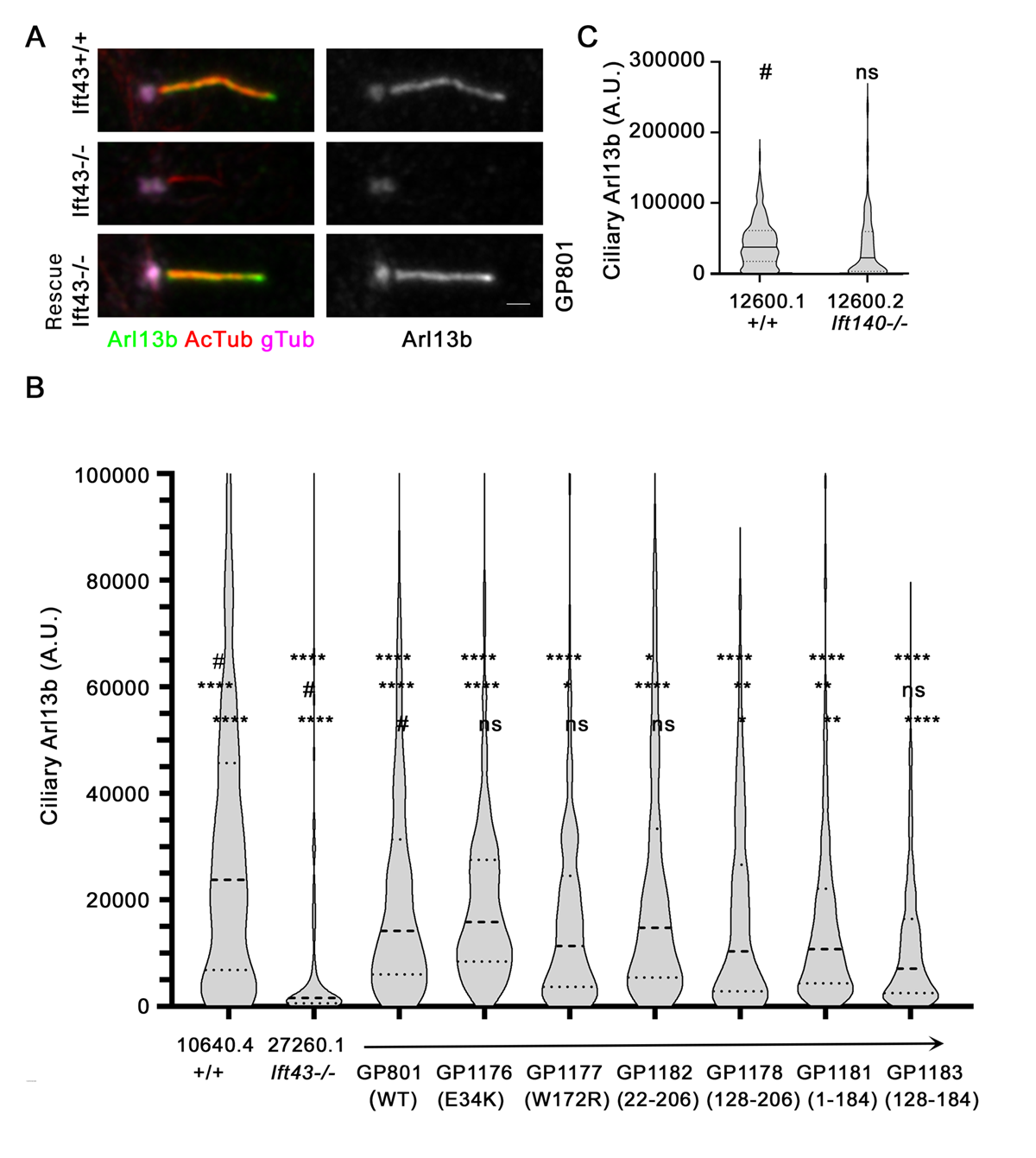
